# The kinetic landscape of an RNA binding protein in cells

**DOI:** 10.1101/2020.05.11.089102

**Authors:** Deepak Sharma, Leah L. Zagore, Matthew M. Brister, Xuan Ye, Carlos E. Crespo-Hernández, Donny D. Licatalosi, Eckhard Jankowsky

## Abstract

Gene expression in higher eukaryotic cells orchestrates interactions between thousands of RNA binding proteins (RBPs) and tens of thousands of RNAs ^1^. The kinetics by which RBPs bind to and dissociate from their RNA sites are critical for the coordination of cellular RNA-protein interactions ^2^. However, these kinetics were experimentally inaccessible in cells. Here we show that time-resolved RNA-protein crosslinking with a pulsed femtosecond UV laser, followed by immunoprecipitation and high throughput sequencing allows the determination of binding and dissociation kinetics of the RBP Dazl for thousands of individual RNA binding sites in cells. This kinetic crosslinking and immunoprecipitation (KIN-CLIP) approach reveals that Dazl resides at individual binding sites only seconds or shorter, while the sites remain Dazl-free markedly longer. The data further indicate that Dazl binds to many RNAs in clusters of multiple proximal sites. The impact of Dazl on mRNA levels and ribosome association correlates with the cumulative probability of Dazl binding in these clusters. Integrating kinetic data with mRNA features quantitatively connects Dazl-RNA binding to Dazl function. Our results show how previously inaccessible, kinetic parameters for RNA-protein interactions in cells can be measured and how these data quantitatively link RBP-RNA binding to cellular RBP function.

---

The binding and dissociation of RBPs to their cognate sites on cellular RNAs are critical for the regulation of gene expression ^2^. Yet, association and dissociation kinetics of RBPs at individual cellular binding sites have not been experimentally accessible. In cells, steady-state patterns of RNA-protein interactions have been measured ^3–5^. RBP binding and dissociation kinetics have only been determined *in vitro* ^2,6^. For a small number of RBPs, equilibrium binding parameters measured *in vitro* correlate with steady-state binding patterns in cells ^7,8^. These observations advanced understanding of RBP function. However, inaccessibility of binding and dissociation kinetics of RBPs in cells severely limits or even precludes the establishment of quantitative connections between RBP-RNA interactions and cellular RBP function. Here, we describe an approach to measure binding and dissociation kinetics of the RBP Dazl at thousands of individual binding sites in cells. We then show how these kinetic parameters inform a quantitative understanding of the cellular function of Dazl.

## Time-resolved fs laser crosslinking

To measure binding and dissociation kinetics of proteins at individual RNA sites in cells, we devised a time-resolved RNA-protein crosslinking approach. (**Fig.1a**). Kinetic parameters in cells have to be determined from the steady-state between free and RNA-bound protein, which can be accomplished if the crosslinking rate constant exceeds dissociation and apparent association constants (**Fig.1a)**. To achieve sufficiently efficient protein-RNA crosslinking, we employed a pulsed femtosecond (fs) UV laser (**Fig.1b**, **Extended Data Fig.1a**). Pulsed UV lasers had been shown to efficiently photo-crosslink proteins to DNA through multi-photonic excitation of the crosslinking species ^9–12^.

**Figure 1.**
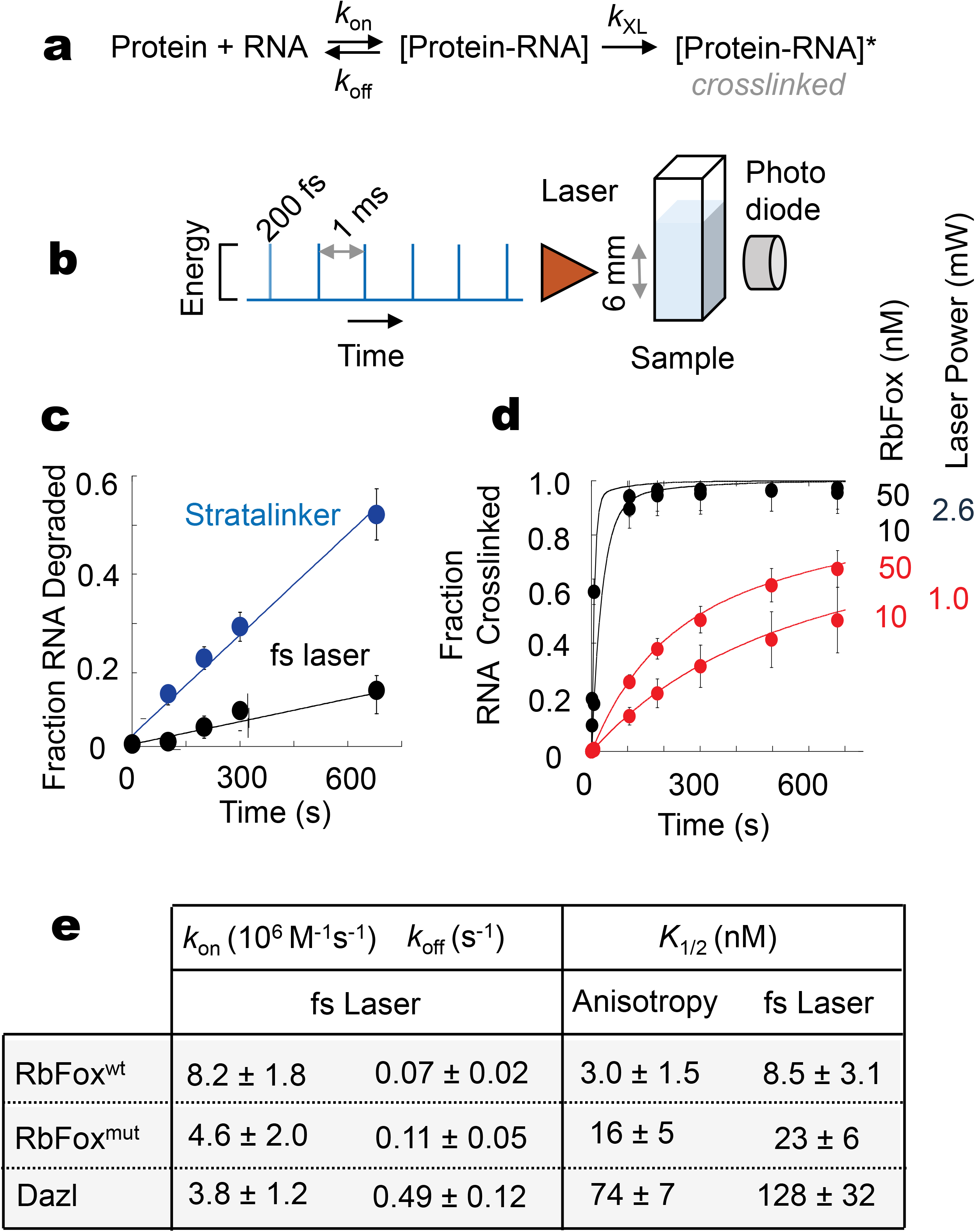
Time-resolved, fs laser RNA-protein crosslinking *in vitro*. **a**. Kinetic scheme for RNA-protein binding and crosslinking. **b**. Schematic of pulsed fs UV laser crosslinking **c**. Degradation of RNA (38 nt) under steady-state and fs laser illumination. Data points represent averages of 3 independent measurements. Error bars mark one standard deviation. Lines show a linear trend. **d**. RNA Crosslinking timecourses for RbFox(RRM) with fs laser at different laser power and protein concentrations. Lines show the fit to the data in panel e. **e**. Association (*k*_on_) and dissociation rate constants (*k*_off_) determined with the fs laser for RbFox(RRM), a mutated RbFox^mut^(RRM), and Dazl(RRM). Equilibrium dissociation constants (*K*_1/2_), calculated from these rate constants and measured by fluorescence anisotropy (**Extended Data Fig.1h-j**). Errors mark one standard deviation.

To examine the utility of a pulsed fs UV laser for determining binding and dissociation rate constants of RNA-protein interactions, we performed time-resolved crosslinking reactions with purified proteins and RNAs (**Fig.1d,e**). UV-mediated RNA degradation was reduced upon irradiation with the fs laser, compared with a steady-state UV light source (**Fig.1c**). Although the photon density during the laser pulse is orders of magnitude greater, compared with the steady-state UV light source, fewer photons are absorbed by the RNA over a given amount of time (**Extended Data Fig.1b**). This is because fs pulses are emitted once per millisecond and the cross-section for multi-photonic absorption is smaller than for single-photonic absorption with a steady-state UV light source ^13^.

Crosslinking of the purified RNA-binding protein RbFox(RRM) to its cognate RNA with the fs laser was markedly more efficient, compared to the steady-state UV source (**Extended Data Fig.1c-e**). Observed crosslinking rates increased with laser power and protein concentration, as expected (**Fig.1d**, **Extended Data Fig.1f,g**). We determined binding and dissociation rate constants for RbFox(RRM)-RNA binding from the crosslinking timecourses at two different laser powers and two different protein concentrations (**Fig.1d,e**). The affinity (*K*_1/2_) of RbFox(RRM) for its cognate RNA, calculated from association and dissociation rate constants was similar to the affinity measured by fluorescence anisotropy (**Fig.1e**, **Extended Data Fig.1h**) and consistent with previously reported values ^14^. We next determined binding and dissociation rate constants for a mutated RbFox^mut^(RRM) ^15^ and for the RNA binding protein Dazl(RRM) ^16^, using fs laser crosslinking (**Fig.1e**). RNA affinities of these two proteins, calculated from the rate constants, were also similar to affinities measured with fluorescence anisotropy (**Fig.1e**, **Extended Data Fig.1i,j**). The data with three RBPs collectively indicate that binding and dissociation rate constants for RNA-protein interactions can be determined by time-resolved, fs laser crosslinking.

## Laser crosslinking in cells

We adapted the time-resolved fs laser crosslinking approach to measure binding and dissociation rate constants of the RNA-binding protein Dazl to individual RNA sites in mouse GC-1 cells ^17,18^. Dazl is essential for male and female gametogenesis ^19–22^. The protein contains one RNA recognition motif (RRM), binds predominantly to 3’UTRs of mRNAs and regulates mRNA stability, translation, or both ^23^. Dazl was expressed under the control of a doxycycline-inducible promotor ^17^. Varying the doxycycline concentration allowed measurements at different Dazl concentrations in GC-1 cells (**Extended Data Fig.2a**). To perform time-resolved fs laser crosslinking experiments, cells were transferred to a quartz cuvette and under constant stirring placed in the laser beam. Crosslinking measurements were performed with GC-1 cells expressing two different Dazl concentrations and two different laser intensities for 30, 180 and 680 s (**Extended Data Fig.2b**). We also measured the bulk degree of crosslinking at each time point (**Extended Data Fig.2c**) and determined transcript levels at each Dazl concentration by RNA-Seq. Approximately 10% of cells showed signs of physical damage after crosslinking, which is comparable to cell damage by conventional steady-state UV-crosslinking (**Supplementary Material Table S4**).

We prepared and sequenced cDNA libraries for each timepoint sample and for controls without crosslinking (**Extended Data Fig.2b**, **Supplementary Material Table S5**, refs.^24,25^). Dazl crosslinking sites with the fs laser were virtually identical to sites identified by conventional steady-state UV-crosslinking with respect to RNA types, location in 3’UTRs and crosslinking site characteristics (**Fig.2a**, **Extended Data Fig.2d-f**, ref.^17^). These data show that fs laser crosslinking maintains the characteristics of crosslink sites seen with steady-state UV-crosslinking. Our kinetic crosslinking and immunoprecipitation approach (KIN-CLIP) thus faithfully maps Dazl binding sites.

**Figure 2.**
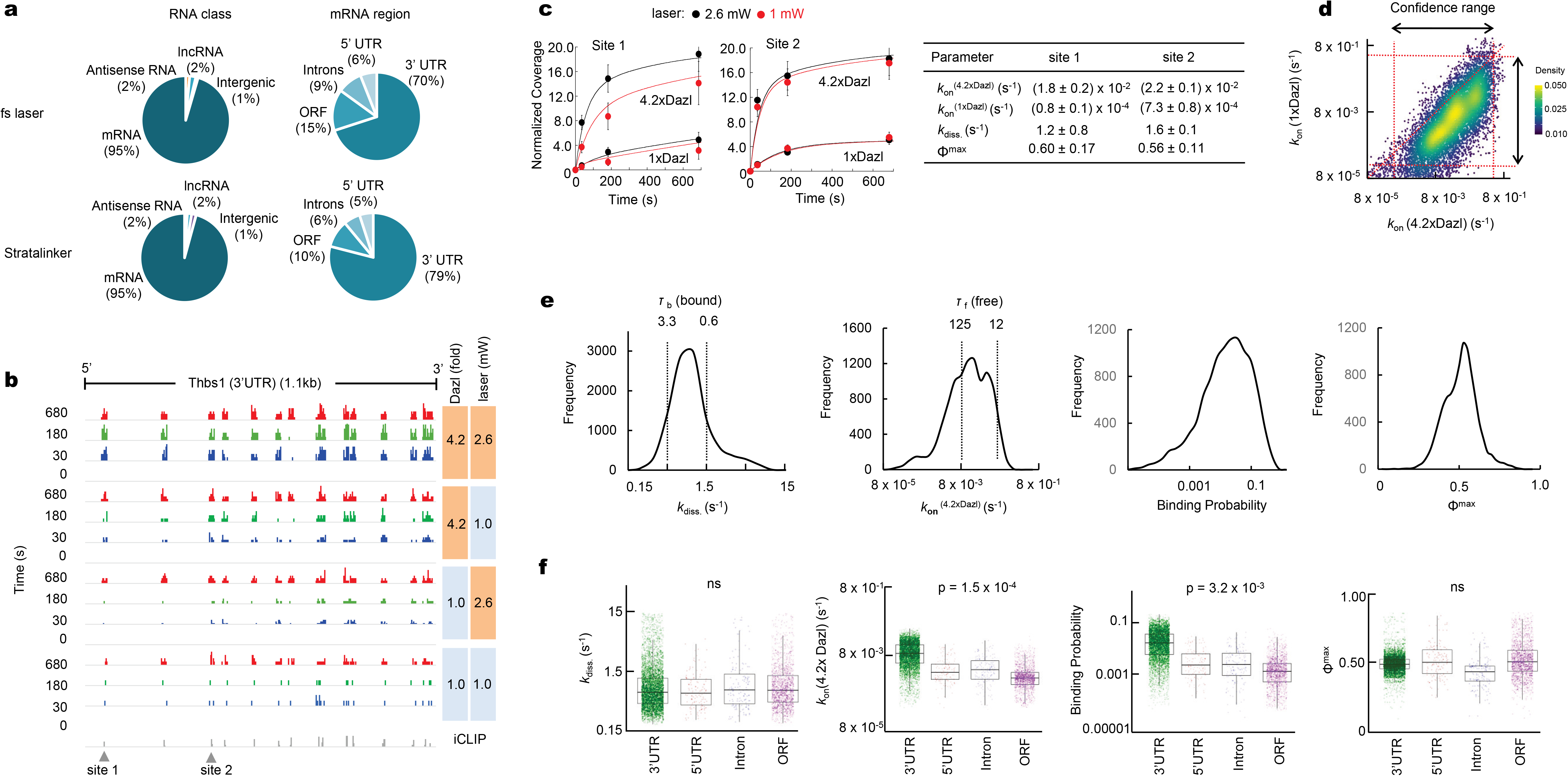
Kinetics of Dazl-RNA binding and dissociation in cells. **a**. Distribution of CLIP sequencing reads across RNA classes and mRNA regions for fs laser (4.2xDazl, 2.6 mW) and conventional crosslinking (Stratalinker; 4.2xDazl). **b**. Normalized sequencing reads for the 3’UTR of a representative transcript (Thbs1) at increasing crosslinking times (left side), different protein concentrations and different laser power (right side). Reads for conventional iCLIP are indicated below. **c**. Crosslinking timecourses for two binding sites (1,2, panel b). Datapoints show the normalized read coverage (lines: best fit to the parameters in the table; error bars: 95% confidence interval for normalized peak coverage value). **d**. Association rate constants for 1xDazl and 4.2xDazl for all binding sites (N = 10,341). Arrows mark the confidence range for the rate constants. The diagonal line marks equal rate constants at both Dazl concentrations. **e**. Transcriptome-wide distributions of dissociation rate constants (*k*_diss._), association rate constants at high Dazl concentration (*k*_on_^4.2xDazl^), binding probability (4.2xDazl), and maximal fractional occupancy (Φ^max^) for all Dazl binding sites. Select dwell times of Dazl bound (*τ*_b_) and away from binding sites (*τ*_f_) are marked. **f**. Distributions of kinetic parameters for all binding sites in the indicated mRNA regions [p-values: One Way ANOVA, n.s.: not significant].

To calculate association and dissociation rate constants for Dazl binding at individual binding sites, we normalized the sequencing reads for each CLIP library to the bulk amount of crosslinking, thereby converting sequencing reads into a concentration-equivalent of crosslinked RNA at a given binding site (**Fig.2b**, **Supplementary Material Table S6**). This normalized read coverage was used to calculate for each binding site a dissociation rate constant (*k*_diss._), observed association rate constants at low and high Dazl concentration (*k*_on_^(1xDazl)^, *k*_on_^(4.2xDazl)^) and crosslinking rate constants for each laser power (*k*_XL_^(1 mW)^, *k*_XL_^(2.6 mW)^, **Fig.2c**, **Extended Data Fig.3a-k**). Obtained rate constants faithfully described the experimental data (**Fig.2c**, **Extended Data Fig.3l,m**).

## Dazl-RNA binding kinetics in cells

For most binding sites (89%), the observed association rate constants at the 1xDazl concentration were lower than those at the 4.2xDazl concentration (**Fig.2d**). These data indicate that only a small fraction of binding sites is saturated with Dazl at low protein concentration and implies a population of free Dazl in the cell, at least at the low Dazl concentration. Although 85% of Dazl crosslinking sites showed the consensus 5’-GUU motif (**Extended Data Fig.4a-d**), association and dissociation rate constants varied by several orders of magnitude (**Fig.2e**). This observation suggests that Dazl binding and dissociation kinetics in cells depend not exclusively on the consensus motif. A_n_, U_n_ and (GU)_n_ stretches were overrepresented in the vicinity of binding sites with high association rate constants (**Extended Data Fig.4e-p**). No further sequence signatures in the vicinity of crosslinking sites correlated with other rate constants (**Extended Data Fig.4i-p**).

The dissociation rate constant for Dazl(RRM) *in vitro* (**Fig.1e**) is on the low end of the spectrum of cellular dissociation rate constants (**Fig.2e**), indicating that Dazl dissociates from most cellular binding sites more frequently than from its cognate RNA *in vitro*. Dazl resides at most cellular binding sites for less than τ_B_ < 1s (**Fig.2e**). Binding events are infrequent and even at high Dazl concentrations occur rarely more than six times per minute (**Fig.2e**). Accordingly, the probability of Dazl to be bound at any time is less than 10% for many binding sites (**Fig.2e**). This observation indicates that Dazl operates at a sub-saturating regime with respect to its mRNA targets in GC-1 cells. This notion is consistent with kinetic parameters of Dazl measured *in vitro* (**Fig.1e**), and a cellular Dazl concentration roughly at or below its affinity *in vitro* ^26^. We also determined a maximal fractional occupancy (Φ^max^, **Fig.2f**), which describes the extent by which a given RNA site would be occupied at saturating Dazl concentrations. The data suggest that most binding sites are not fully accessible for Dazl binding during the course of the experiment.

Dissociation rate constants for binding sites did not vary significantly for different RNA classes (**Extended Data Fig.4s**) or between mRNA 3’UTRs, 5’UTRs, introns and open reading frames (**Fig.2f**). Association rate constants and binding probabilities, which depend on both, association and dissociation rate constants, were higher for binding sites in 3’UTRs than for sites in 5’ UTRs, introns and ORFs (**Fig.2f**), and higher in mRNAs, compared with other RNA classes (**Extended Data Fig.4q,r**). The maximal fractional occupancy of binding sites did not significantly vary in the different mRNA regions (**Fig.2f**), but was higher in mRNA, compared with other RNA classes (**Extended Data Fig.4t**). Since Dazl function has been linked to binding in 3’UTRs ^17^, our data raised the possibility that association rate constants, binding probabilities, or both, influence cellular roles of Dazl more than its residence time at the binding sites. Collectively, the kinetic data revealed highly dynamic Dazl-RNA interactions with most Dazl binding events being rare and transient.

## Dazl binds mRNAs in clusters

To understand how Dazl regulates mRNA function in this highly dynamic fashion, we examined the patterns of the kinetic parameters for all Dazl binding sites on each bound mRNA. The majority of Dazl binding sites are in 3’UTRs (**Fig.2a**), and frequently proximal to the polyadenylation site (PAS, **Extended Data Figs.2e**, **5a**). Most Dazl-bound mRNAs contained multiple Dazl binding sites with an inter-site distance markedly smaller than expected by chance (**Fig.3a**), even when distant to the PAS (**Extended Data Figs.5b,c**). This observation suggested clustering of multiple Dazl binding sites on most 3’UTRs (**Extended Data Fig.5d-g**). The number of binding sites within a 3’UTR cluster increased with proximity to the PAS (**Fig.3b**). Association rate constants for individual binding sites scaled with the number of binding sites in a cluster, regardless of the distance of the cluster to the PAS. (**Fig.3c**). Binding probabilities showed a similar pattern (**Extended Data Fig.5h**). Dissociation rate constants and maximal fractional occupancies did not scale with the number of binding sites in a cluster (**Extended Data Fig.5i,k**). Kinetic parameters within clusters showed consistent patterns of moderate correlation (**Extended Data Fig.5k**). Fractional occupancies for binding sites within a given cluster were closely correlated (**Fig.3d**, **Extended Data Fig.5k**), suggesting a similar accessibility of binding sites within a cluster. This notion, together with the scaling of association rate constants with the number of binding sites (**Fig.3c**), raised the possibility that binding site clusters are important for Dazl function.

**Figure 3.**
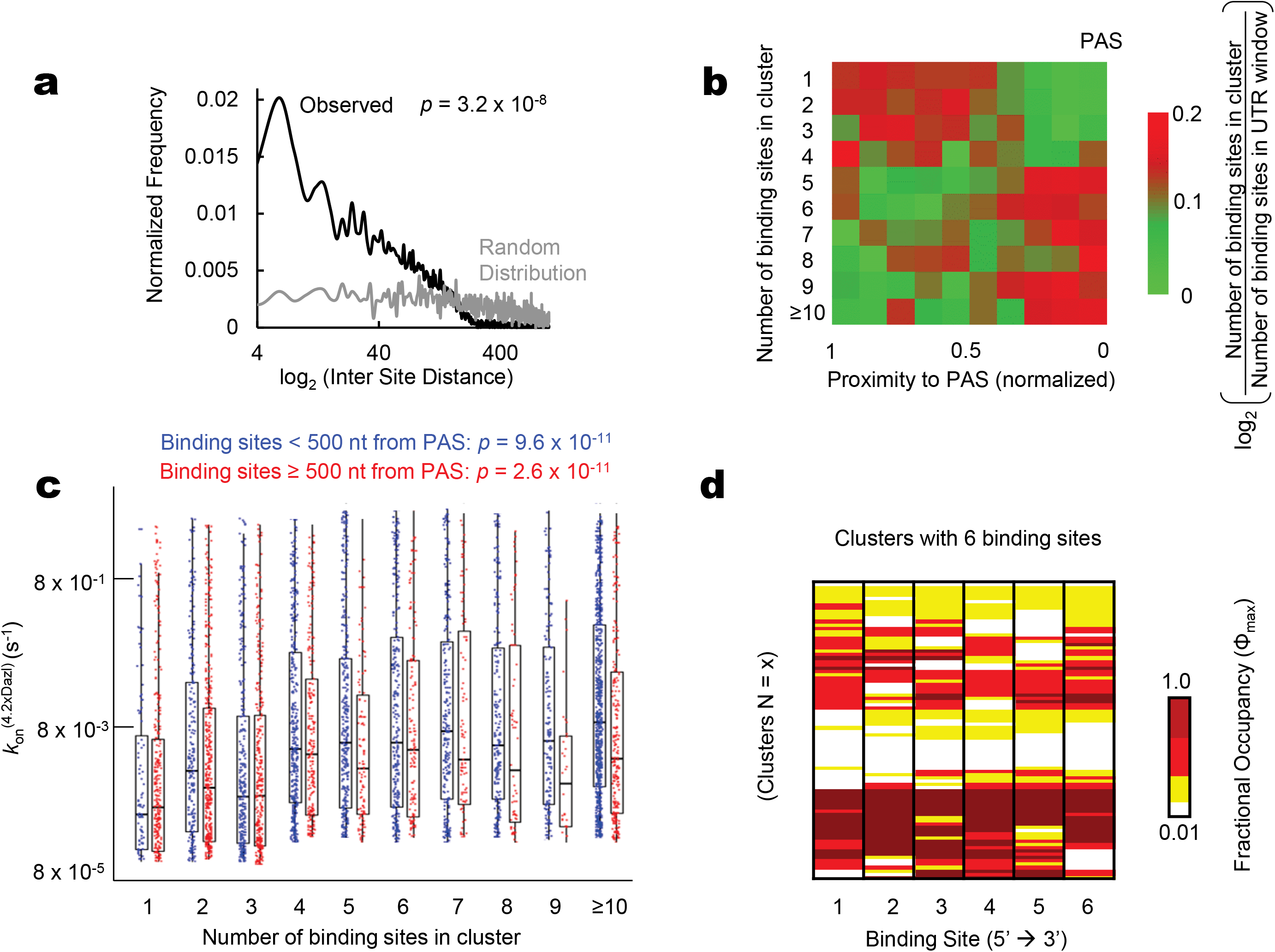
Clustering of Dazl binding sites in 3’UTRs. **a**. Distribution of Dazl binding sites in 3’UTRs as function of the distance between neighboring binding sites. The grey line shows the distribution if sites were randomly distributed across all 3’UTRs (*p* value: t-test). **b**. Proximity of clusters with varying number of binding sites to the PAS. **c**. Correlation between association rate constants and number of binding sites in clusters. (p-values: one way ANOVA). **d**. Heatmap depicting correlation of values for maximal fractional occupancy in clusters with 6 binding sites.

## Clusters correlate with Dazl function

To test this hypothesis, we quantified Dazl binding in a given cluster by calculating a cumulative binding probability (ΣB) from the kinetic constants of the binding sites in the cluster. ΣB describes the probability that Dazl occupies at least one site in a given cluster at any given time (**Fig.4a**). ΣB increased with the number of binding sites in a cluster and with proximity to the PAS (**Extended Data Fig.6a,b**). We compared ΣB values in a given cluster to changes in ribosome association and transcript levels at low and high Dazl concentrations (**Fig.4b**). Dazl binding had been shown to increase transcript levels and ribosome association for many, but not all mRNAs ^17^. We detected an overrepresentation of clusters with high ΣB in mRNAs that increased in transcript level, ribosome association, or both, at the high Dazl concentration, compared with the low Dazl concentration (**Fig.4c**, **Extended Data Fig.6c,d**). Clusters with low ΣB values were overrepresented in mRNAs that decreased in transcript levels and ribosome association at the high Dazl concentration (**Fig.4c**). We detected no comparable correlation between the Dazl impact on transcript levels or ribosome association and binding probabilities of individual binding sites, clusters with scrambled binding sites or with simultaneous occupancy of multiple binding sites in a given cluster (**Extended Data Fig.6e-k**). ΣB values thus instructively link binding kinetics to Dazl impact on mRNA function, further supporting the notion that Dazl clusters are critical for the function of this RBP.

**Figure 4.**
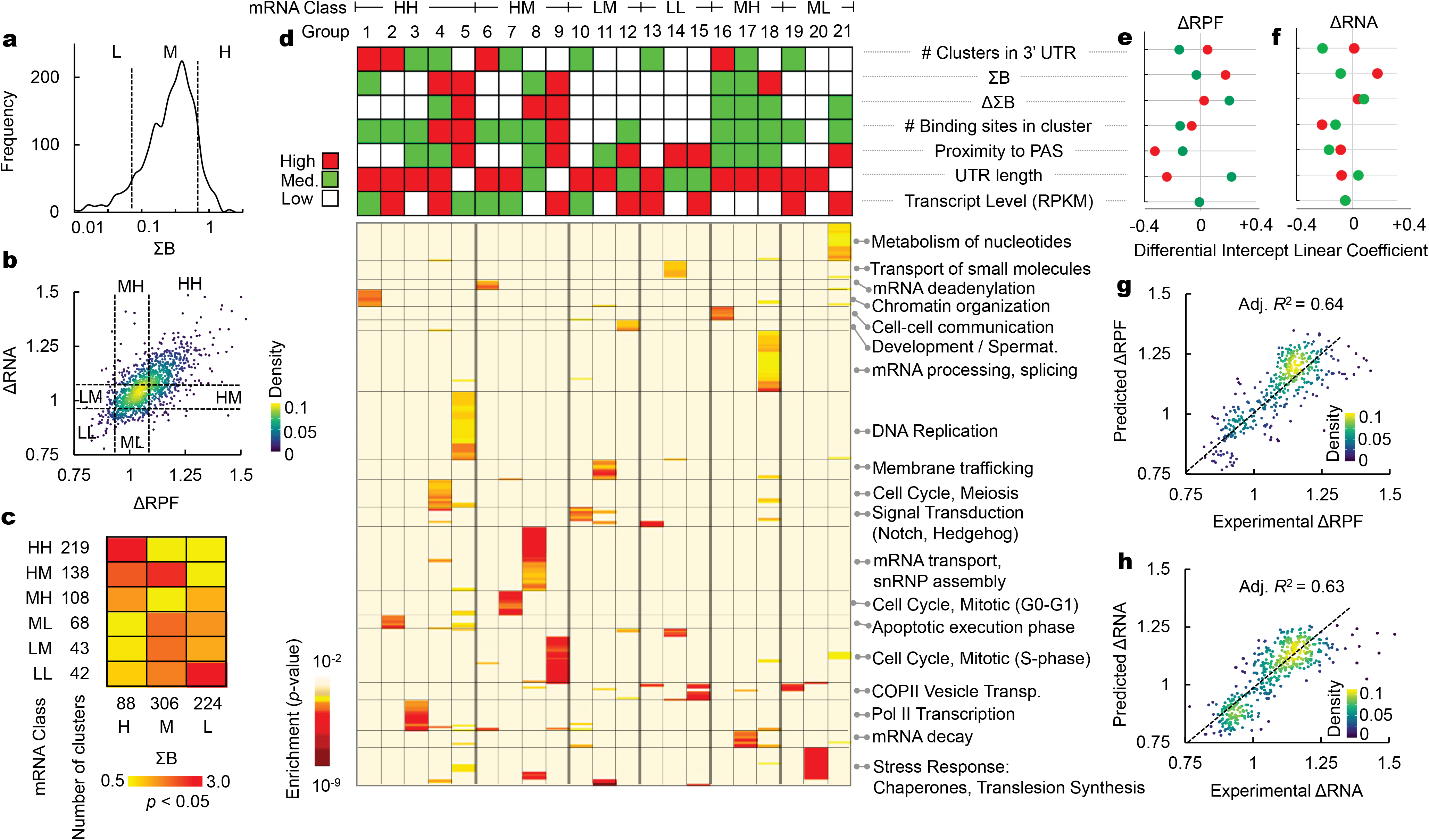
Link between Dazl-RNA binding and Dazl impact on mRNA function. **a**. Distribution of cumulative binding probabilities (ΣB) for Dazl in all clusters (N = 1,690). **b**. Changes in transcript levels (ΔRNA) and ribosome association (ΔRPF) between 4.2xDazl and 1xDazl for Dazl-bound mRNAs (N = 968). Colors mark the functional mRNA classes (first letter; ΔRPF, second letter ΔRNA; L-low, M-medium, H-high). Data points represent averages from triplicate ribosome profiling and RNAseq experiments ^17^. **c**. Correlation between cumulative binding probabilities and functional mRNA classes. Colors correspond to the enrichment (hypergeometric test, red: *p* < 0.05, shades of yellow: not enriched) **d**. Upper panel: Heatmap of Dazl-code, linking functional mRNA classes to kinetic parameters (ΣB, ΔΣB), cluster characteristics (number of binding sites in cluster, cluster distance from PAS) and UTR features (numbers of clusters, on UTR, UTR length, transcript level), all shown in terciles (**Extended Data Fig.8f**). Numbers mark the groups with characteristic combinations of ΣB, ΔΣB, cluster and mRNA features. Lower panel: Link between Dazl-code and Gene ontology (GO) terms. **e,f**. Linear regression model linking Dazl-code to impact of Dazl binding on changes in transcript levels (ΔRNA) and ribosome association (ΔRPF) (panel b). Points represent the differential intercept (DI) linear coefficient (LC) (red: DILCs for transcript levels and ribosome association that increase at high Dazl concentration, green: DILCs for transcript levels and ribosome association that decrease at high Dazl concentration). **g,h**. Correlation between experimental values for ΔRNA and ΔRPF (N = 434) and values calculated with the linear regression model (R: adjusted linear correlation coefficient).

## The Dazl code

To delineate the connection between Dazl binding kinetics and Dazl impact on mRNA function in more detail, we identified additional mRNA and Dazl cluster characteristics that correlated with Dazl function. Besides ΣB, we detected correlations for the number of binding sites in a cluster, the difference in cumulative binding probabilities at low and high Dazl concentrations (ΔΣB), number of clusters in a 3’UTR, length of the 3’UTR, and proximity of a cluster to the PAS (**Extended Data Fig.7**). Some of these characteristics correlate with each other (R^2^ ≤ 0.6), but each parameter contributes separately to the Dazl impact on mRNA function (**Extended Data Fig.8a-e**). Proximity of Dazl binding to the PAS had been previously noted to influence Dazl impact on mRNA function ^17^.

Principal component analysis and t-distributed stochastic neighbor embedding independently identified 21 mRNA groups with a distinct combination of kinetic, cluster and mRNA characteristics (**Extended Data Fig.8b-e**). Each of these 21 groups falls into a class of Dazl impact on transcript level and ribosome association (**Fig.4d**, **Extended Data Fig.8c-f**). The mRNAs in each group belong to defined GO-terms (**Fig.4d**), and in many cases encode proximal proteins in a given pathway (**Extended Data Fig.8h**). These findings indicate a link between the biological role of a given mRNA and Dazl binding kinetics, binding site clusters, their location on the 3’UTR and mRNA features (**Extended Data Figs.8h**,**9**). Collectively, these parameters represent a basic Dazl code that connects Dazl binding in 3’UTRs to its impact on mRNA function (**Fig.4d**, **Extended Data Fig.8f**)

To quantify this Dazl code, we employed a linear regression model (**Fig.4e,f**; **Extended Data Fig.10**), which explains roughly 64% of experimental variance in the scaling of transcript level and ribosome association with the Dazl concentration (**Fig.4g,h**). The largest contribution is seen for the cumulative binding probabilities, which derive from the kinetic parameters of Dazl binding, and for the numbers of Dazl clusters in the 3’UTR (**Fig.4e,f**). For mRNAs that increase in ribosome association, the distance of the Dazl clusters to the PAS also has an effect (**Fig.4e**), consistent with the previously reported data ^17^. Collectively, our data show that Dazl impacts bound mRNAs in a complex, yet tractable manner that depends prominently on kinetic parameters.

## Discussion

We devised and applied a time-resolved crosslinking approach that provides transcriptome-wide access to cellular binding and dissociation kinetics of RNA-protein interactions at individual binding sites. Key to this KIN-CLIP approach is a pulsed fs UV laser, which increases crosslinking efficiencies without altering RNA-protein crosslinking patterns, compared with steady-state UV irradiation. KIN-CLIP should enable the biochemical characterization of other RNA-protein interactions in cells. Combining time-resolved fs laser crosslinking and kinetic analysis might also allow quantitative, biochemical analysis of DNA-protein ^12^ and even of protein-protein interactions ^27^ in cells.

For Dazl, KIN-CLIP reveals highly dynamic RNA binding. Dazl resides at individual binding sites for only seconds or shorter, while cognate sites remain free of Dazl for most of the time. These findings are consistent with kinetic data for Dazl-RNA binding *in vitro* and the notion that cellular Dazl concentrations are sub-saturating relative to its RNA targets ^26^. Highly dynamic binding allows for rapid changes in RNA binding patterns, which might be critical for Dazl function. *In vitro* RNA binding kinetics of Dazl are similar to those of other RBPs ^6^, many of which might also operate in cells at sub-saturating concentrations relative to their RNA targets ^7^. Our findings raise the possibility that other RBPs bind their cognate RNA sites also transiently and infrequently. If true for many RBPs, few, if any regulatory RBPs might be bound to a given mRNA at any given time.

Access to cellular kinetic data allows the decoding of a complex link between Dazl-RNA-binding patterns and Dazl function. Dazl regulates mRNA level and ribosome association according to a code that integrates the collective binding kinetics of Dazl at its cognate sites in a cluster, the number of binding sites in a cluster, location of clusters on the 3’UTR, proximity to the PAS, and 3’UTR length. Since our experimental and data analysis approaches are applicable to other RBPs, KIN-CLIP provides a blueprint for delineating other RBP-codes.

## Acknowledgements

We thank Dr. Gabriele Varani (UW, Seattle) for the gift of purified RbFox(RRM) and RbFox^mut^(RRM), Dr. Anton Komar (Cleveland State University) for the design of the codon-optimized Dazl construct, and Dr. Wei Huang (CWRU) for assistance with the fluorescence polarization experiments. This work was supported by the NIH (GM118088 to E.J., GM107331 to D.D.L) and the NSF (CHE-1800052 to C.E.C.-H.)

## EXTENDED DATA, FIGURE CAPTIONS

**Extended Data Figure 1.**
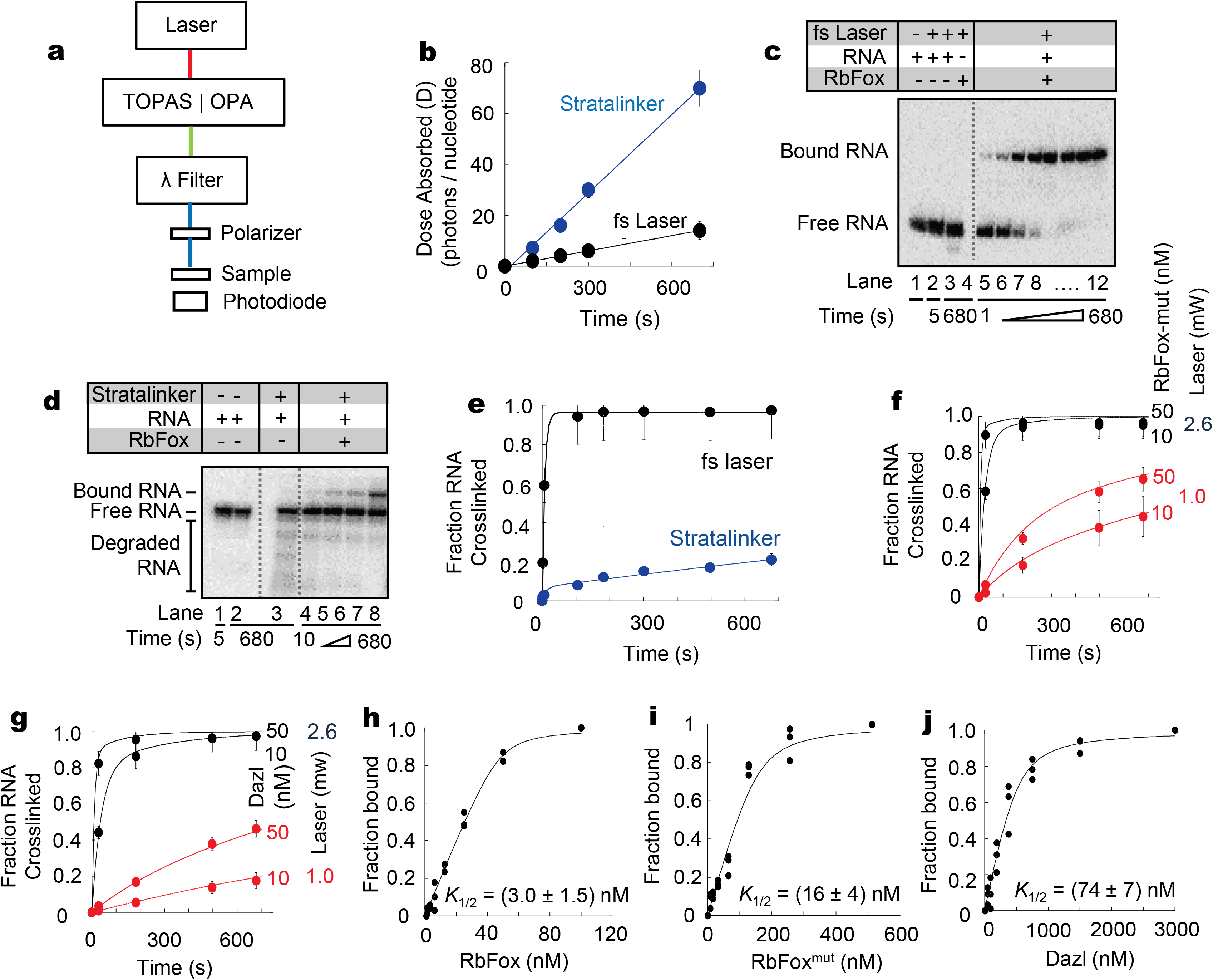
Time-resolved RNA-protein crosslinking with fs laser *in vitro*. **a**. Schematics of fs laser setup. **b**. Dose absorbed over time for crosslinking with conventional UV (Stratalinker, 200 mJ/cm^2^, λ = 254 nm) and fs laser (2.6 mW) **c**. Representative denaturing <u>p</u>oly<u>a</u>crylamide <u>g</u>el <u>e</u>lectropherogram (PAGE) for a crosslinking reaction of 50 nM RbFox(RRM) (laser: 2.6 mW) (lanes 5 – 12) and control reactions with RNA only (lanes 1 – 3) and RbFox(RRM) only (lane 4), with (lanes 2-4) or without (lanes 1 and 5) crosslinking. **d**. Representative denaturing PAGE for a crosslinking reaction of 50 nM RbFox(RRM) with Stratalinker (200 mJ/cm^2^, λ = 254 nm), lanes 4 – 8) and control reactions (lanes 1 – 3). **e**. Timecourse of crosslinking reaction of 50 nM RbFox(RRM) with Stratalinker (200 mJ/cm^2^, λ = 254 nm) vs. fs laser (**Fig.1d**). Datapoints are averages from triplicate experiments (error bars: one standard deviation). **f**. RNA Crosslinking timecourses for RbFox^mut^(RRM) with fs laser at different laser power and protein concentrations. Data points represent averages of 3 independent measurements (error bars: one standard deviation). Lines show the fit to the data in **Fig.1e**. **g**. RNA Crosslinking timecourses for Dazl(RRM) with fs laser at different laser power and protein concentrations. Data points represent averages of 3 independent measurements (error bars: one standard deviation). Lines show the fit to the data in **Fig.1e**. **h-j**. Binding isotherms for RbFox(RRM), RbFox^mut^(RRM) and Dazl(RRM) to cognate RNAs measured by fluorescence anisotropy. Experiments were performed multiple times, all datapoints are shown. Apparent equilibrium binding constants (*K*_1/2_, **Fig.1e**) were calculated with the quadratic binding equation.

**Extended Data Figure 2.**
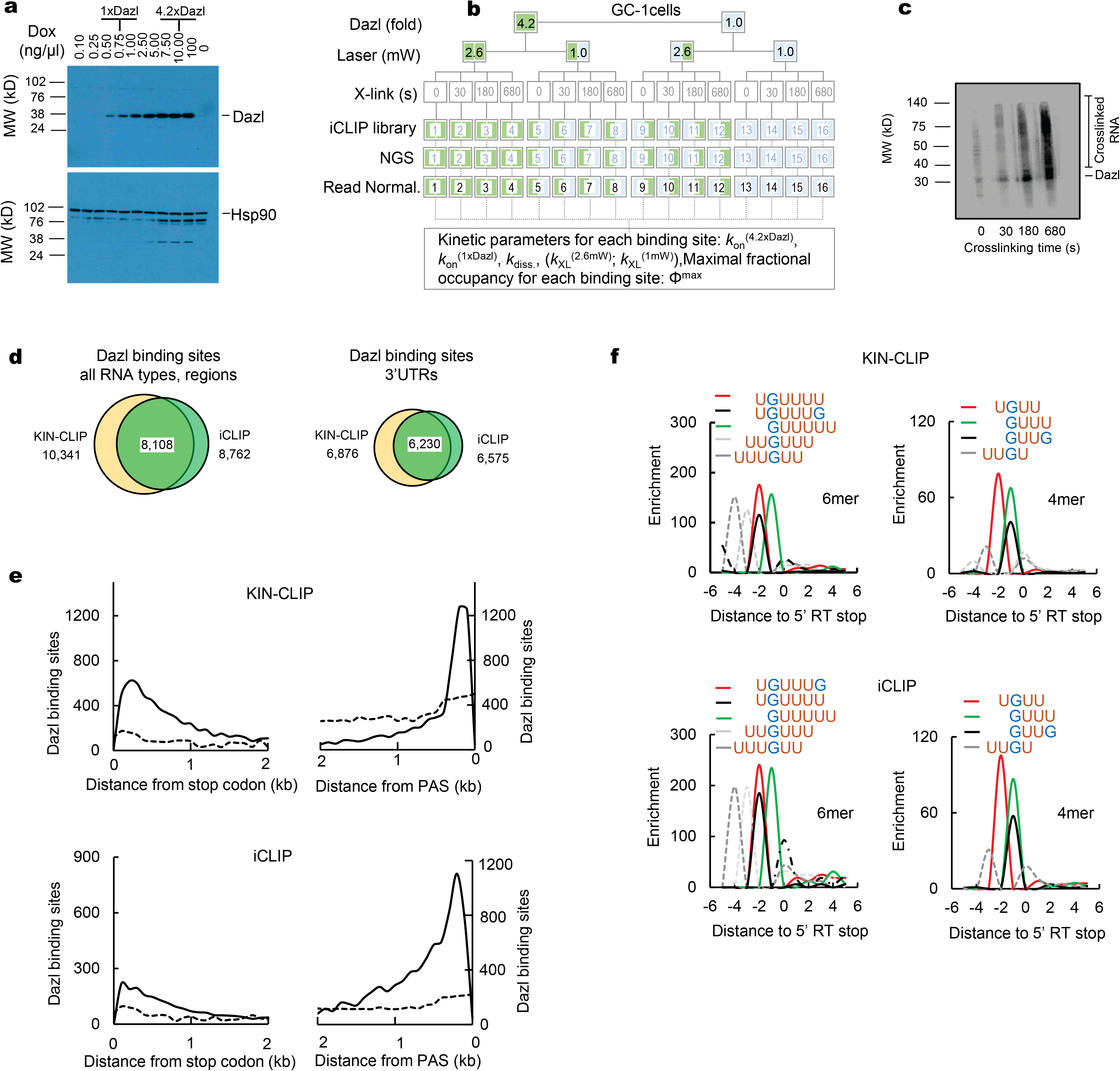
Dazl-RNA crosslinking with fs laser in GC-1*spg* cells. **a.** Western Blot of Doxycyline dependent Dazl expression in GC-1 cells. **b**. Schematic of the time-resolved crosslinking approach in cells. Numbers mark the respective CLIP libraries. **c**. Representative PAGE for bulk Dazl-RNA crosslinking. The intensity of crosslinked RNA (marked) is used to convert NGS reads to a concentration-equivalent parameter (for bulk crosslinking intensities see **Supplementary Material, Table S6**) **d**. Dazl binding sites identified by fs laser (KIN-CLIP) and conventional UV crosslinking (iCLIP) on all RNAs and 3’UTRs. **e**. Metagene distribution of Dazl binding sites identified by KIN-CLIP and iCLIP on 3’UTRs proximal to stop codon and PAS. The dotted lines mark the background of a random distribution of binding sites on 3’UTRs. **f**. CITS (<u>Cr</u>osslink <u>I</u>nduced <u>T</u>runcation <u>S</u>ite) analysis ^28,29^ of 6-mer and 4-mer enrichment at 5’-termini of sequencing reads for KIN-CLIP (upper panels) and iCLIP (lower panels). The data indicate a virtually identical sequence context of crosslinking sites for KIN-CLIP and iCLIP. Sequence enrichment reflects the statistical overrepresentation of 6-mer and 4-mer sequences with respect to randomized sequences (Z-score, 11 nucleotide region, ± 5 nt from the 5’-terminal nucleotide).

**Extended Data Figure 3.**
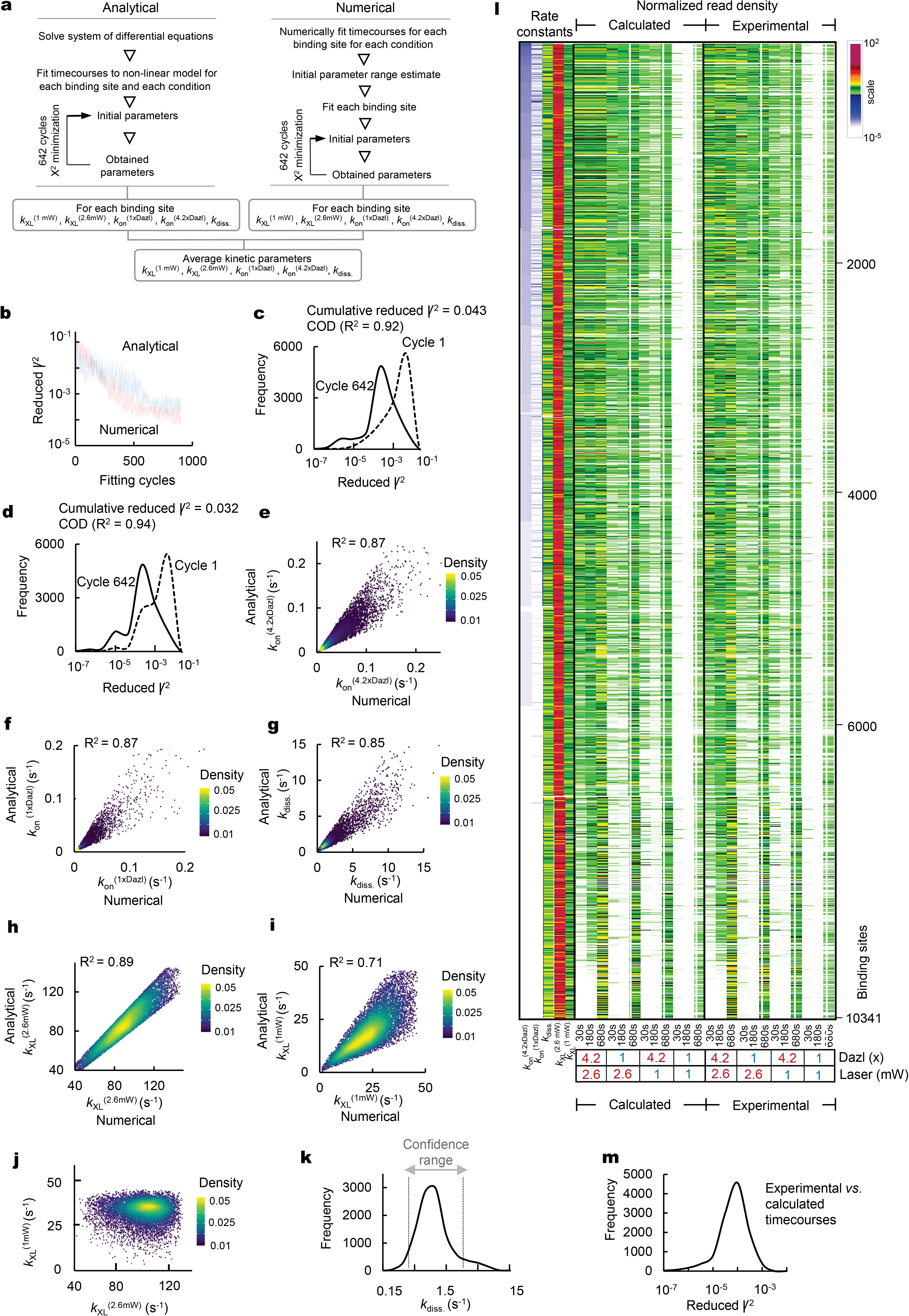
Determination of kinetic parameters from fs laser, time-resolved Dazl-RNA crosslinking in cells. **a**. Flowchart of the approach to calculate kinetic parameters for individual Dazl-RNA binding sites in cells (for details see Materials and Methods). Unless otherwise stated, rate constants averaged from both approaches are used in subsequent data analyses. **b**. Scaling of Χ^2^ with the number of iterative fitting cycles for analytical and numerical approaches. **c,d**. Distribution of Χ^2^ at first and last (642) fitting cycle for analytical (**c**) and numerical (**d**) approaches (COD: Coefficient Of Determination, *R*^2^: linear correlation coefficient). **e-i**. Correlation of parameters calculated with analytical and numerical fitting procedures (*R*^2^: linear correlation coefficient). **j**. Correlation between crosslinking rate constants for low and high laser power. Rate constants are averaged from parameters obtained with numerical and analytical approach. Crosslinking rate constants at higher laser power were larger than at lower for 92% of binding sites. **k**. Confidence range for dissociation rate constants (for details see Materials and Methods). **l**. Normalized read densities measured experimentally and calculated from the kinetic parameters for all Dazl binding sites. **m.** Distribution of Χ^2^ for experimental values compared with values calculated with the kinetic parameters.

**Extended Data Figure 4.**
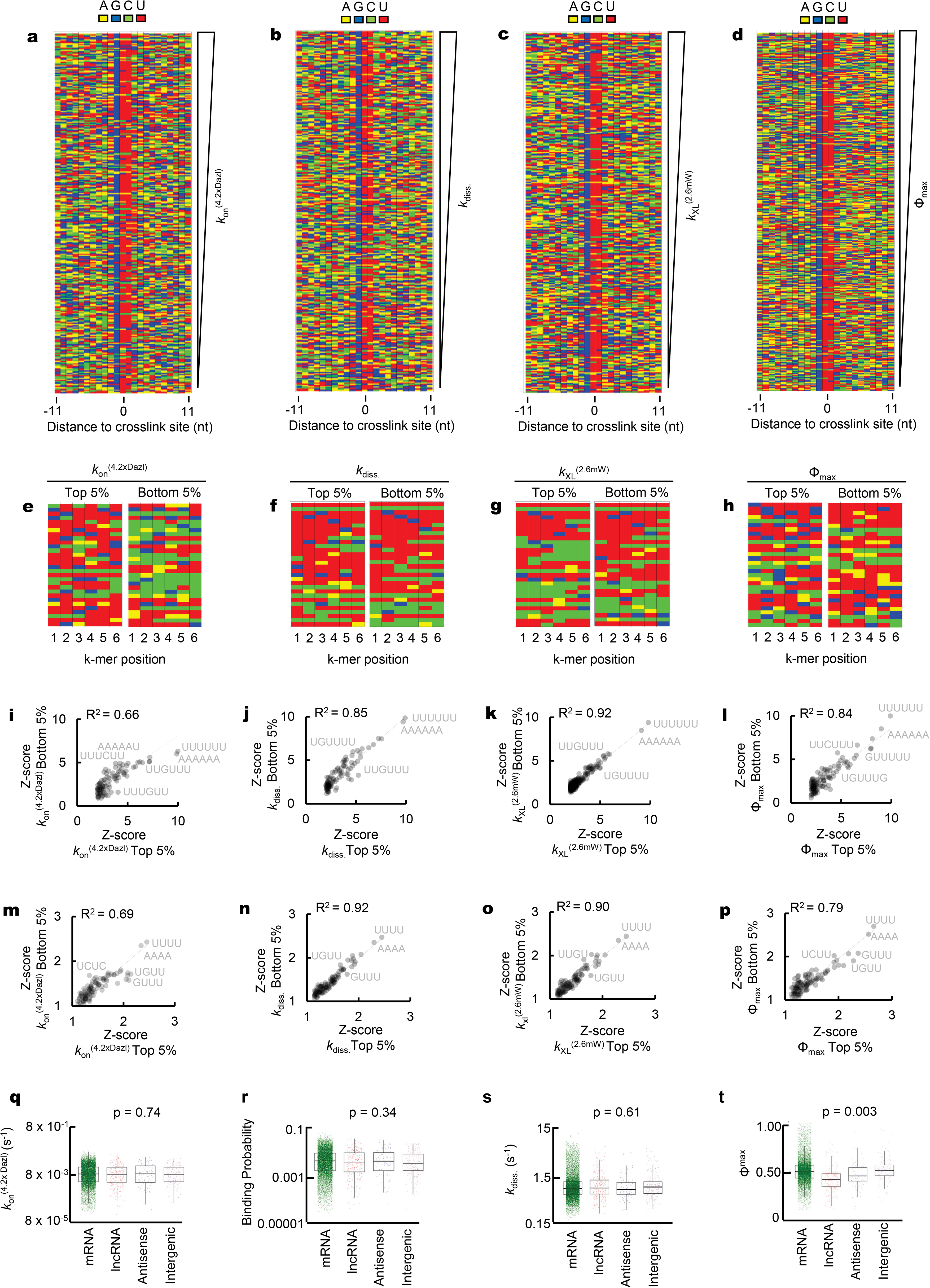
Kinetic parameters of Dazl binding sites and sequence context. **a-d**. Sequences surrounding Dazl binding sites, arranged according to decreasing values for *k*_on_^(4.2xDazl)^, *k*_diss._, *k*_XL_^(2.6mW)^, and Φ^max^. Sequences are aligned at the peak nucleotide (most frequent crosslink site, **Extended Data Fig.2f**, position 0). **e-h**. Frequency of 6-mer sequences surrounding Dazl crosslink sites (± 11 nt peak nucleotide) in top and bottom 5% of sequences arranged according to the kinetic parameters in panels (**a-d**). **i-l**. Relative frequency of 6-mer sequences in top and bottom 5% of sequences (panels **e-h**), arranged according to the kinetic parameters in panels **a-d**. Sequences below the diagonal line correspond to enrichment of a 6-mer in the top 5% versus the bottom 5%. (*R*^2^: linear correlation coefficient). A_6_, U_6_ and U_3_GU_2_ are most enriched in the vicinity of the binding sites with the fastest apparent association rate constants, compared to the binding sites with the slowest apparent association rate constants. No comparable enrichment is seen for other kinetic parameters. **m-p**. Relative frequency of 4-mers in top and bottom 5% of sequences arranged according to the kinetic parameters in panels (**a-d**). **q-t**. Distribution of association and dissociation rate constants, binding probabilities (P) and maximal fractional occupancy (Φ^max^) for binding sites on different RNA classes. P values (one-way ANOVA, significant for p < 0.05) indicate inter-group differences. Φ^max^, but not other parameters vary significantly for different RNA classes.

**Extended Data Figure 5.**
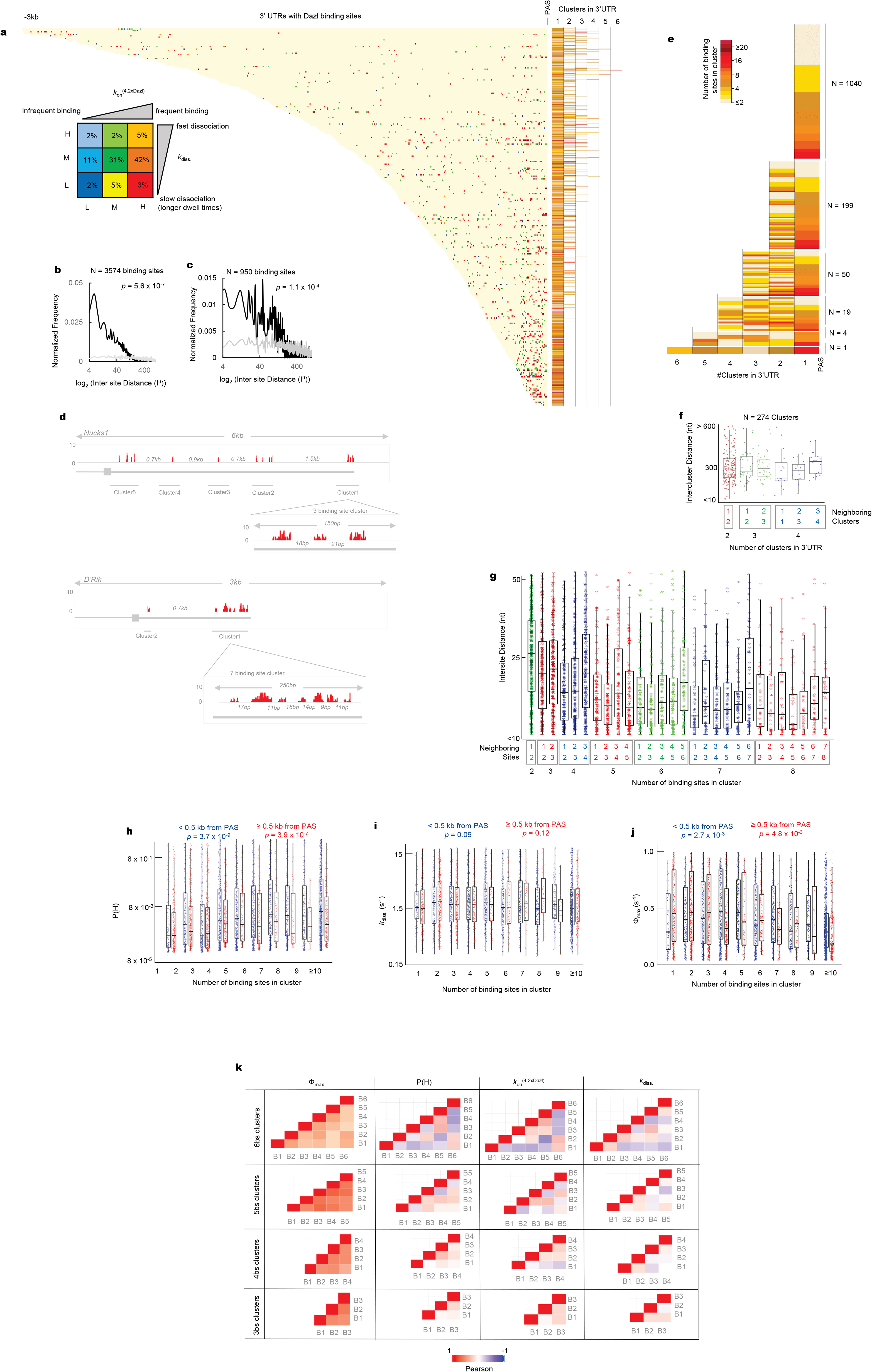
Arrangement of 3’UTR Dazl binding sites in clusters. **a**. Arrangement of Dazl binding sites in 3’UTRs. Binding sites are colored according to *k*_on_^(4.2xDazl)^ and *k*_diss._ as indicated in the key panel. Right panel: number of clusters in corresponding 3’UTR. Colors mark number of binding sites in a cluster, as indicated in legend bar (right) (N = 1,313 3’UTRs, 1,690 clusters, 6,085 binding sites) **b**. Distribution of Dazl binding sites in 3’UTRs closer than 500 nt to PAS, as function of the distance between neighboring binding sites. The grey line shows the distribution if sites were randomly distributed across all 3’UTRs (*p* value: t-test). **c**. Distribution of Dazl binding sites in 3’UTRs farther than 500 nt from PAS, as function of the distance between neighboring binding sites. The grey line shows the distribution if sites were randomly distributed across all 3’UTRs (*p* value: t-test). **d**. Large windows: genome browser traces of representative 3’UTRs with 5 clusters (Nucks1) and 2 clusters (D’Rik, D030056L22Rik). Bars show the normalized read coverage for 4.2xDazl, 2.6 mW laser and 680s crosslinking time. Numbers mark the distance between clusters. Small windows: zoom into cluster 1 of Nucks1 with 3 binding sites and in cluster 1 of D’Rik with 2 binding sites (numbers mark the distance between binding sites). **e.** Number of clusters in 3’UTRs with Dazl binding sites. Colors show the number of binding sites in a cluster as indicated in panel **a**. (red: 20; cornsilk: 1). **f**. Distances between clusters in 3’UTRs with 2 to 4 clusters. Number 1 represents the cluster most proximal to the PAS. **g**. Distribution of distances between neighboring binding sites in clusters (2-9 binding sites). Number 1 represents the 3’ binding site. **h-j**. Correlation between the number of binding sites for clusters proximal (blue: < 0.5 kb) and distant (red: ≥ 0.5 kb) to the PAS and (*P*^(4.2xDazl)^, **h**), dissociation rate constants (*k*_diss._, **i**), and maximal fractional occupancy (Φ^max^, **j**), for individual binding sites in a given cluster. P-values (one way ANOVA) indicate significant inter-group differences for *P*^(4.2xDazl)^ and Φ^max^, but not for *k*_diss._ *P*^(4.2xDazl)^ and Φ^max^ depend on *k*_on_^(4.2xDazl)^, which correlates with the number of binding sites in a cluster, (**Fig.3c**). **k**. Correlation between kinetic parameters of individual binding sites in clusters with 6, 5, 4, and 3 binding sites. The Pearson correlation coefficient is indicated in the legend bar. Binding site number 1 indicates the 3’ binding site in a cluster.

**Extended Data Figure 6.**
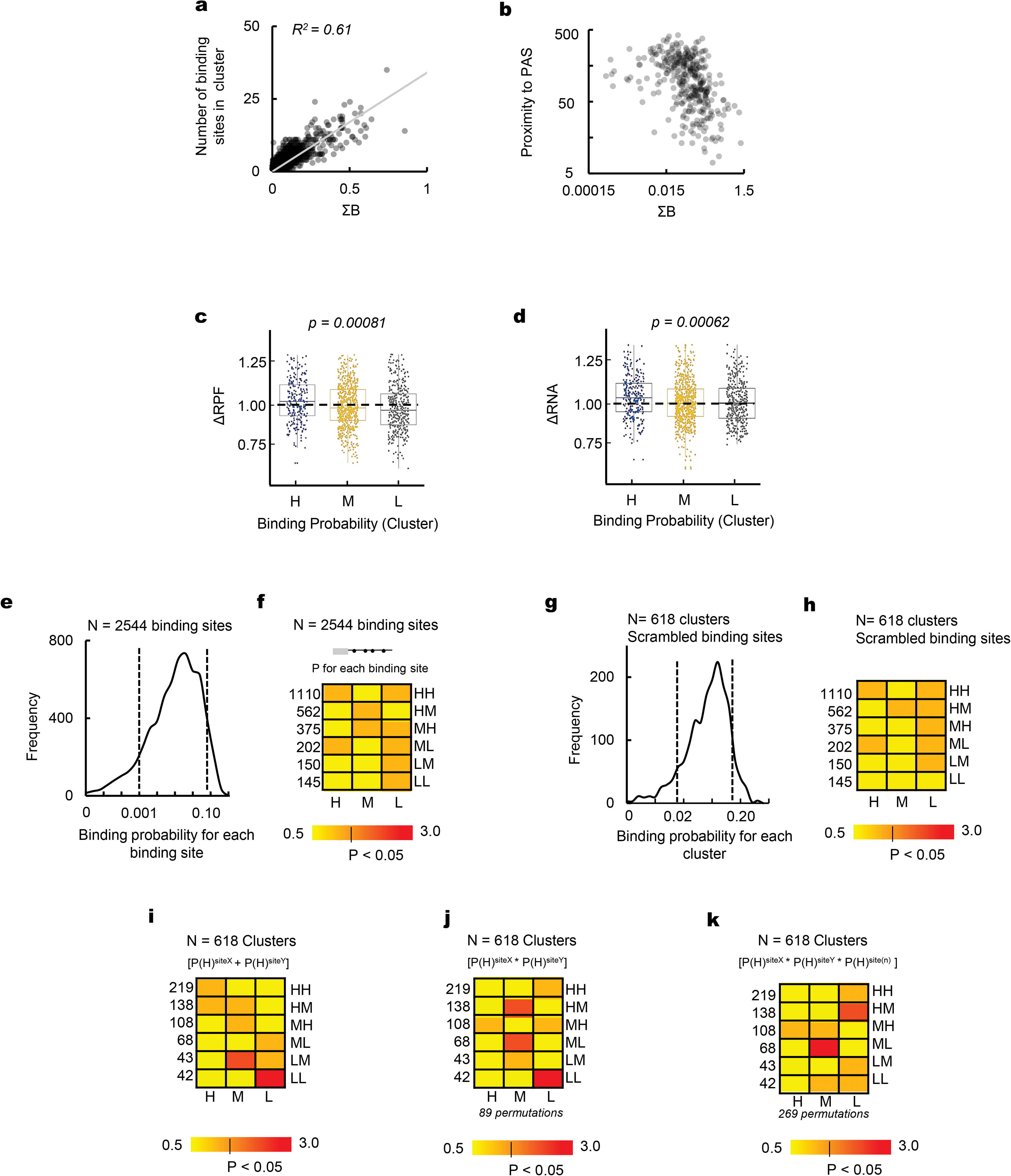
Link between Dazl binding in 3’UTRs and impact on mRNA level and ribosome association. **a**. Correlation between cumulative binding probabilities (ΣB) and number of binding sites in a cluster (N = 1,313 3’UTRs, 6,085 binding sites, 1,690 clusters), *R*^2^: linear correlation coefficient). **b**. Correlation between ΣB and distance of the cluster from the PAS, *R*^2^: linear correlation coefficient). **c**. Correlation of ΣB terciles (H: high; M: medium; L: low, **Fig.4a**) and changes in ribosome association (ΔRPF, **Fig.4b**) for the corresponding transcripts (N = 968) between low (1xDazl) and high (4.2xDazl) concentration (P value: one-way ANOVA). For UTRs with multiple clusters, the cluster closest to the PAS was utilized. **d**. Correlation of ΣB terciles (H: high; M: medium; L: low, **Fig.4a**) and changes in transcript levels (ΔRNA, **Fig.4b**) for the corresponding transcripts between low (1xDazl) and high (4.2xDazl) concentration (P value: one-way ANOVA). For UTRs with multiple clusters, the cluster closest to the PAS was utilized. **e**. Distribution of binding probabilities for individual Dazl binding sites in 3’UTRs for transcripts in HH, HM, MH, LM, LL and ML mRNA classes (**Fig.4b**). The dotted lines mark terciles (H: high; M: medium; L: low), (for details, see Materials and Methods). **f**. Correlation between binding probabilities for individual binding sites and functional mRNA classes (**Fig.4b**). Colors mark the enrichment (hypergeometric test, red: p < 0.05, shades of yellow: not enriched, see color bar). No significant enrichment is observed. **g**. Distribution of cumulative binding probabilities for Dazl clusters in 3’UTRs with scrambled binding sites. The dotted lines mark terciles (H: high; M: medium; L: low). **h**. Correlation between cumulative binding probabilities of Dazl clusters with scrambled binding sites (panel **g**) and functional mRNA classes (**Fig.4b**). Colors mark the enrichment (hypergeometric test, Red: p < 0.05, shades of yellow: not enriched, see color bar). No significant enrichment is observed. **i.** Correlation between additive binding probabilities of two Dazl sites in a cluster and functional mRNA classes (hypergeometric test, red: p < 0.05, shades of yellow: not enriched, see color bar). For clusters with > 2 binding sites, permutations of two sites were tested and sites with highest additive binding probability were selected. The model tests whether the additive binding probability of any two Dazl binding sites in a given cluster can explain the impact of Dazl on the transcript to the same extent as considering cumulative binding probabilities for the entire cluster (**Fig.4c**). The model is only able to explain the LL, LM mRNA classes, which frequently contain transcripts with clusters that have only few Dazl binding sites. **j**. Correlation between conditional binding probabilities of two Dazl sites in a cluster (terciles) and functional mRNA classes (hypergeometric test, Red: p < 0.05, shades of yellow: not enriched, see color bar). For clusters with > 2 binding sites, permutations of two sites were tested and combinations of sites with the highest multiplicative binding probability were selected. The model tests whether the conditional binding probability of any two Dazl binding sites (e.g. whether Dazl needs to bind simultanously to both sites) in a given cluster can explain the impact of Dazl on the transcript to the same extent as considering cumulative binding probabilities for the entire cluster (**Fig.4c**). The model explains only mRNA classes which frequently contain transcripts with Dazl clusters that have only few binding sites. For these clusters cumulative and conditional binding probabilities scale similarly. The data suggest that simultaneous binding of Dazl to two sites in a cluster is not required for general Dazl function. **k**. Correlation between conditional binding probabilities of three Dazl sites in a cluster (terciles) and functional mRNA classes (hypergeometric test, Red: p < 0.05, shades of yellow: not enriched, see color bar). For clusters with > 3 binding sites, permutations of three sites were tested and combinations of sites with the highest multiplicative binding probability were selected. The model tests whether the conditional binding probability of three Dazl binding sites (e.g. whether Dazl needs to bind simultanously to three sites) in a given cluster can explain the impact of Dazl on the transcript to the same extent as considering cumulative binding probabilities for the entire cluster (**Fig.4c**). The model explains only mRNA classes which frequently contain transcripts with Dazl clusters that have only few binding sites. For these clusters cumulative and conditional binding probabilities scale similarly. The data suggest that simultaneous binding of Dazl to two or more sites in a cluster is not required for Dazl function.

**Extended Data Figure 7.**
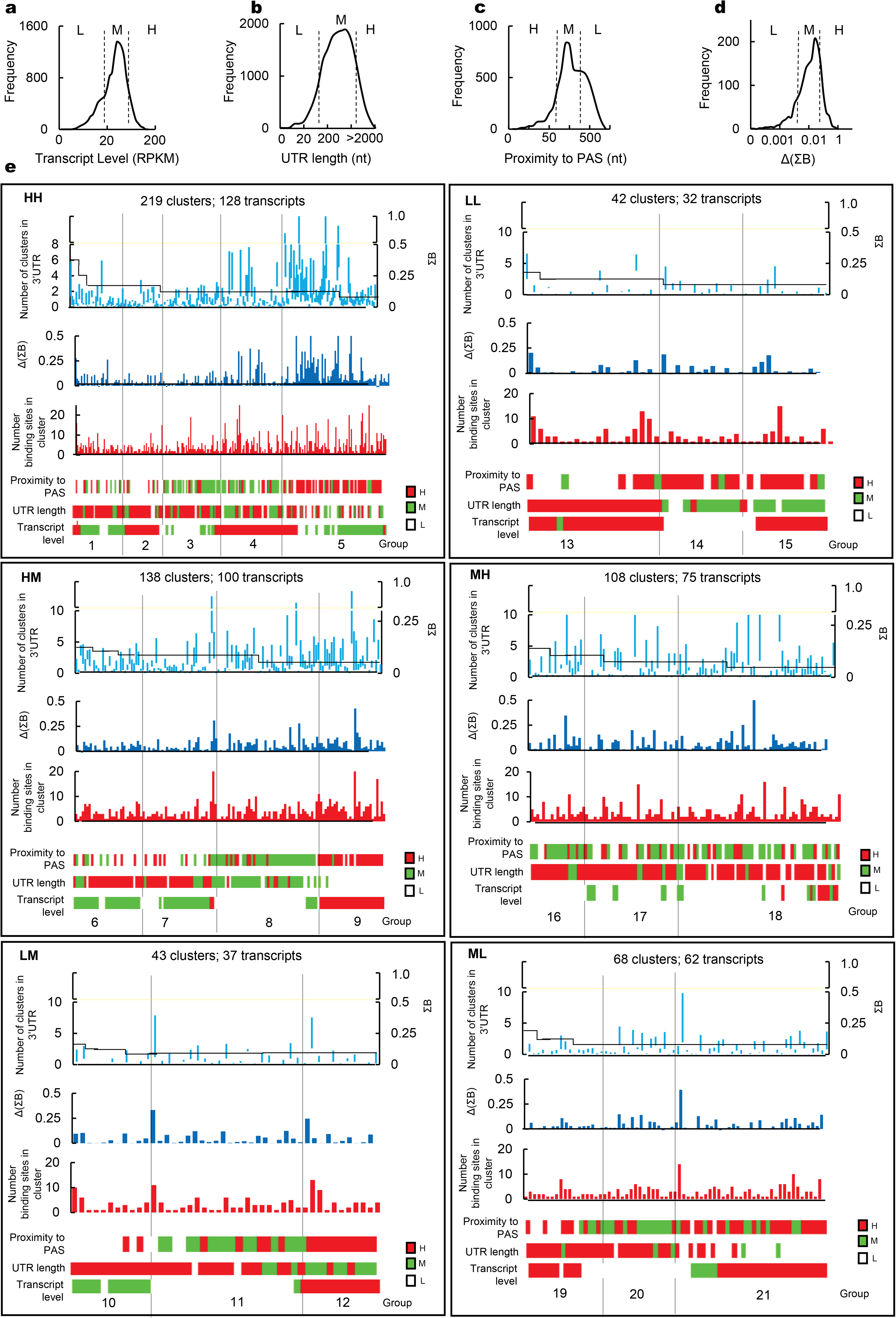
Link between Dazl clusters in 3’UTRs and impact on mRNA level and ribosome association. **a.** Distribution of transcript levels at 4.2xDazl **b.** Distribution of 3’UTR lengths ^17,30,31^. For UTRs with multiple lengths, coordinates for the longest 3’UTR were utilized. **c.** Distribution of distances of Dazl clusters from PAS. **d.** Distribution of differential cumulative binding probability (ΔΣB) for all Dazl clusters. The dotted lines mark terciles (H: high; M: medium; L: low). Terciles were defined by obtained standard deviations from the mean for each feature described above. **e**. Link between Dazl impact on mRNA level and ribosome association and cluster features (upper graphs: number of Dazl clusters in 3’UTR: black line; ΣB: blue vertical lines, lower end marking ΣB at 1 x Dazl, upper end ΣB at 4.2 × Dazl; middle graphs: ΔΣB for each cluster and number of Dazl binding sites in each cluster; Heatmaps below the graphs: terciles of transcript features obtained from panels **a**-**c**. Each panel shows one functional mRNA class (defined in **Fig.4b**; first letter: change in ribosome association, second letter change in transcript level upon increase in Dazl concentration. H-high (increase at high Dazl concentration), M-medium (no change), L-low (decrease at high Dazl concentration)]. Functional classes not displayed contained too few or no transcripts (LH: 0, HL: 2) or showed no change in ribosome association and transcript level (MM).

**Extended Data Figure 8.**
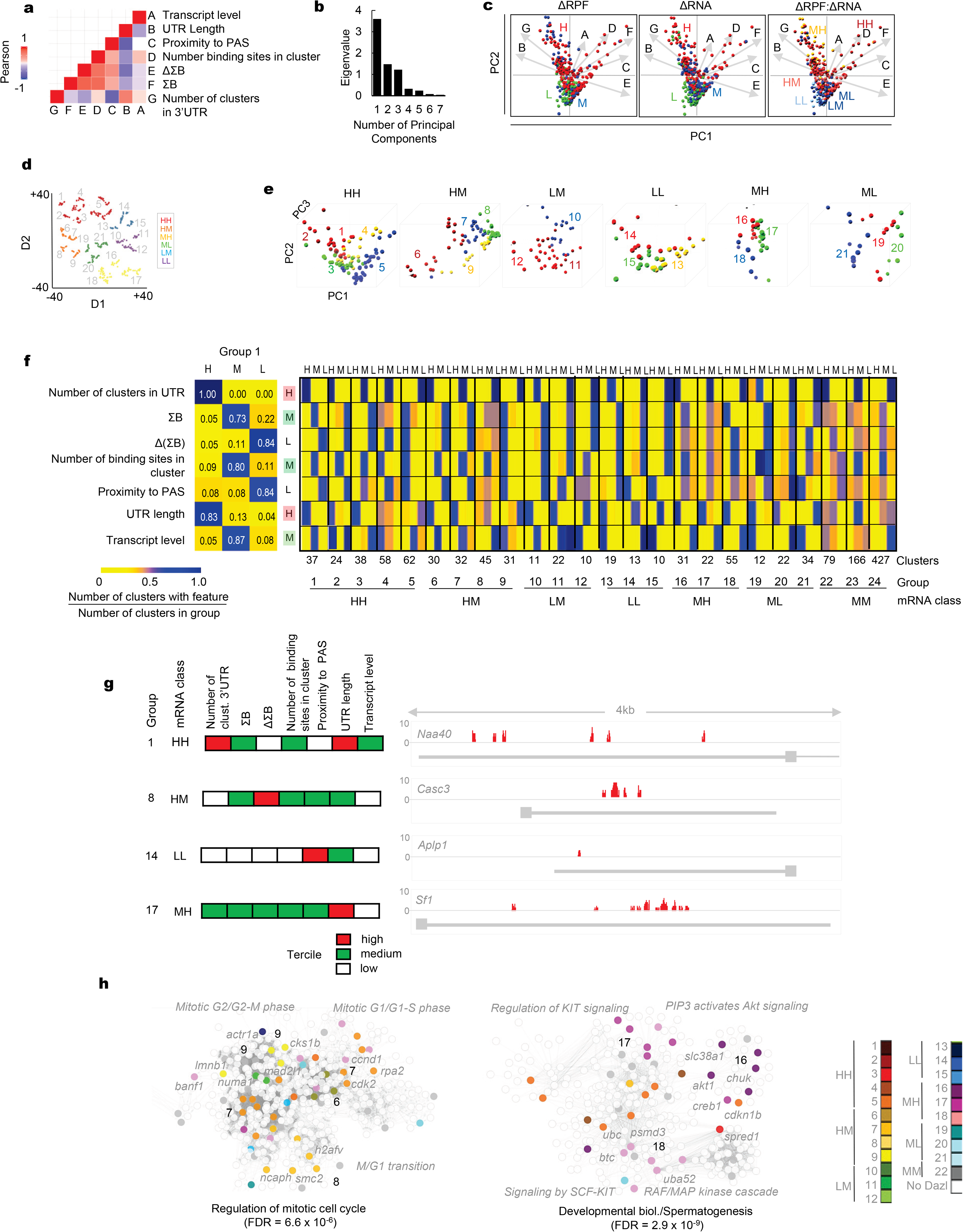
The Dazl code. **a**. Pairwise correlation between Dazl cluster features. Colors correspond to Pearson’s’ correlation coefficient. Cluster features are marked as indicated on the right. **b**. Variance of data reflected in the eigenvalues of principal component axes (N = 7) obtained by PCA. Each eigenvalue corresponds to a principal component axis. Each axis reflects a linear combination of Dazl cluster features (N = 7, obtained from panel (**a**). The eigenvalues and the corresponding principal component axis are sorted by according to the initial variance they represent. The first three principal component axes explain roughly 90% variance. **c**. Biplots of Dazl cluster features (arrows) projected on the first two principal components (PC1,2; panel **b**). Dots represent transcripts. Colors correspond to terciles of the distributions of values for ΔRPF (H = High, M: Medium, L: Low, **Fig.4b**), ΔRNA ((H = High, M: Medium, L: Low, **Fig.4b**), and functional mRNA classes (HH, HM, LM, LL, MH and ML, **Fig.4b**). Each arrow represents a cluster feature (labels as in panel (**a**)). Proximity of arrows scales with correlation between the corresponding features. Arrows in the x-direction (positive or negative) contribute to PC1, arrows in the y-direction (positive or negative) contribute to PC2. Short arrows (transcript level, proximity to PAS) indicate that additional principal components (PC3-7) are required to explain the corresponding feature. **d.** T-distributed Stochastic Neighbor Embedding (t-SNE, Perplexity = 10, Iterations = 2,000) of cluster features (panel **a**). Identified groups are marked 1-21. Each point represents a transcript. **e.** Biplots of Dazl cluster features (arrows) projected on three principal components (PC1,2,3, panel **b**). Dots represent transcripts. Colors correspond to functional mRNA classes (HH, HM, LM, LL, MH and ML, **Fig.4b**). Separation of transcripts in 21 groups is marked as 1-21. **f**. Link of functional mRNA classes to kinetic parameters (ΣB, ΔΣB), cluster features (number of binding sites in cluster, proximity to PAS) and UTR features (numbers of clusters on UTR, UTR length, transcript level). Left panel: enrichment of terciles (H, M, L; **Fig.4a**, **Extended Data Fig.7a-d**) for ΣB, ΔΣB, number of binding sites in cluster, cluster distance from PAS, UTR length and transcript level in group 1. Numbers and color indicate the degree of enrichment. The row on the right marks the visualization of the Dazl code for group 1 that is used in **Fig.4d**. Right panel: enrichment of terciles for the features indicated in the left panel for all groups (1-21). Functional mRNA classes for the respective groups are shown on the bottom. **g**. Genome browser traces of representative transcripts of select groups. mRNA classes are indicated. The y-axis represents normalized coverage value. **h**. Mapping of transcripts from select groups on two biological networks. Groups are colored as indicated in the legend. Proximity of transcripts of a given group in the network indicates closely related biological functions.

**Extended Data Figure 9.**
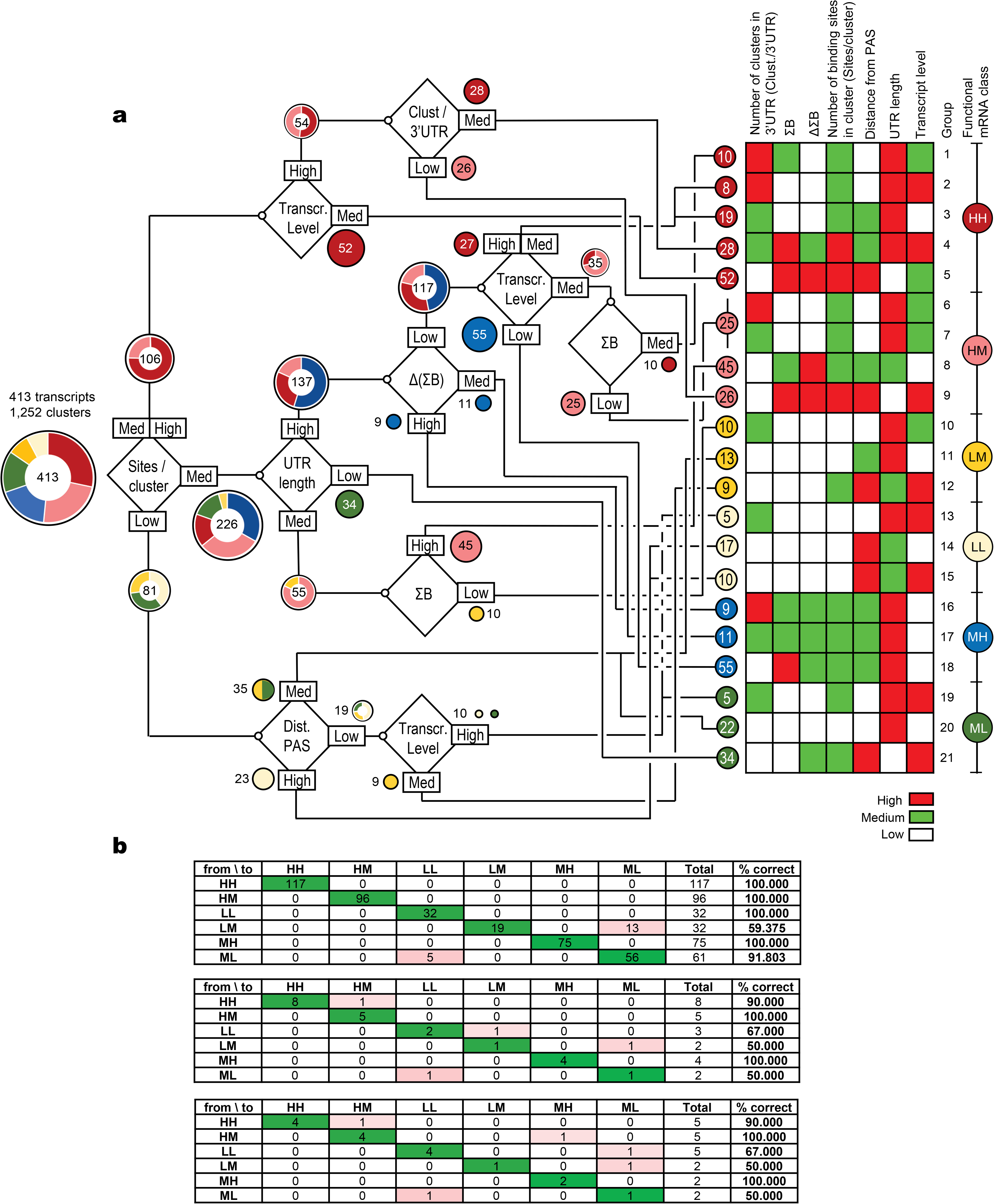
Decision tree classification linking the Dazl code to functional impact of Dazl binding. **a.** Decision tree classifier (Chi-squared automatic interaction detection (CHAID) algorithm ^32–34^ of 7 features (ΣB, ΔΣB, distance to PAS, 3’UTR length, transcript level; Clust/UTR: number of clusters in a given 3’UTR, **Extended Data Fig.8**) in terciles (H: high, M: medium, L: low, **Extended Data Fig.7**). Nodes (◊) mark the given feature and corresponding partition (high, medium, low). Circles indicate the number of transcripts, donut graphs mark the functional mRNA classes, color coded as shown on the right. Circled numbers left to the heatmap with the Dazl code (identical to that in **Fig.4d**) indicate the number of transcripts in a given group. The decision tree was calculated by cross-tabulation of predictor variables (transcripts, N = 413) with target variables (functional mRNA classes HH, HM, MH, LM, LL, ML, **Fig.4b**) followed by partitioning of predictor variables into statistically significant subgroups (Χ^2^ test, for independence with significance threshold: 0.05 (ref.^35^). **b**. Confusion matrix indicating corresponding to the decision tree. Validation 1 (N = 24 transcripts) and Validation 2 (N = 21 transcripts) are predictions for transcripts that were not included in the decision tree classification.

**Extended Data Figure 10.**
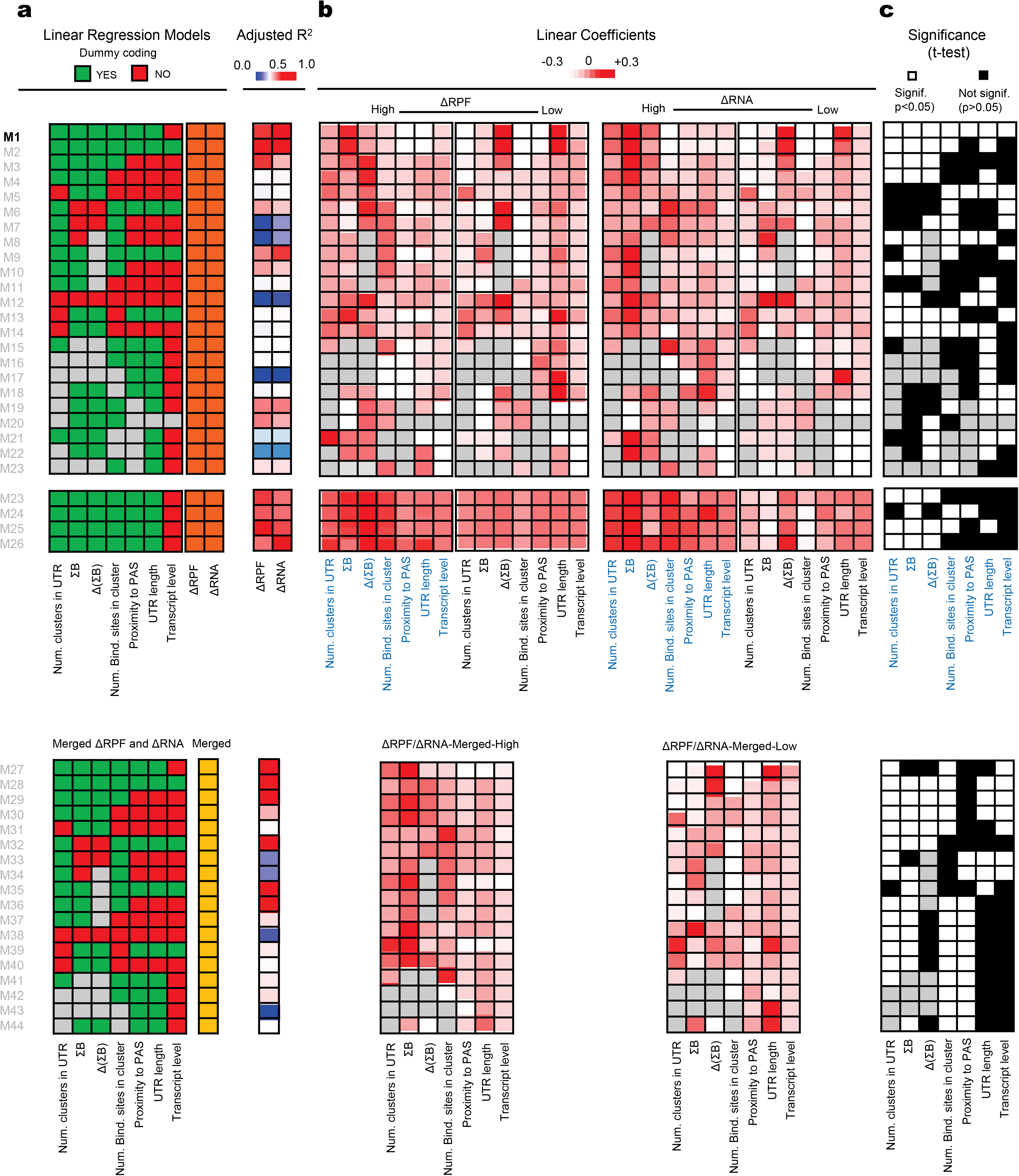
Linear regression models for linking the Dazl code to Dazl impact on changes in transcript levels and ribosome association. **a**. Linear Regression models tested. (Green: dummy coding, using terciles of the variables, **Extended Data Fig.8**. Red: no dummy coding; use of continuous data. Grey: variable was omitted. Yellow: merged data for ΔRPF and ΔRNA). Right columns: adjusted *R*^2^ for each model. **b**. Differential Intercept Linear Coefficients (DILC) for each model. Grey boxes mark models without the respective variable. **c**. Significance of each DILC for each model (White: p < 0.05 – significant, Black p > 0.05 – not significant, p-values: student t-test on each coefficient term). M1 is the only model with consistently significant DILCs.

## MATERIALS AND METHODS

### Laser Setup

The cross-linking experiments were performed by using a Ti:Sapphire regenerative amplifier laser system (Libra-HE, Coherent, Inc.; λ = 800 nm (center wavelength, nominal), pulse width ≤100 fs (Full Width at Half Maximum), 4.0 W at 1 kHz, contrast ratio > 1000:1 pre-pulse; > 100:1 post-pulse; root mean square (8 h) energy stability under stable environmental conditions after system warmup < 0.5 %). The 800 nm fundamental beam was converted to the 270 nm excitation beam by second harmonic sum frequency generation with an optical parametric amplifier (TOPAS, Quantronix/Light Conversion)^36,37^. Contributions to the excitation beam from other wavelengths were removed by a set of dichroic mirrors (λ-filter) and a Glan-Taylor polarizer ^37^. The excitation beam was collimated to a spot size of 6.0 mm. The photon flux at the sample was 1.25·10^16^ cm^−2^s^−1^ (2.6 mW) and 4.81·10^15^ cm^−2^s^−1^ (1 mW) at 270 nm with a pulse duration of 200 (± 50) fs, assuming a Gaussian-shaped pulse ^38^. Stability of the laser output at λ = 270 nm was monitored with a silicon photodiode (S120VC, ThorLabs). The power of the excitation beam was attenuated with a neutral density filter for the crosslinking experiments with the average power of 2.6 mW and 1.0 mW. The crosslinking experiments were conducted in a 2 mm optical path length quartz cell with a maximum sample volume of 0.7 mL, placed orthogonal to the excitation beam. Homogeneity of the sample in the cuvette was maintained with a Teflon-coated magnetic stirring bar (Sterna Cells, Inc.) throughout the measurement. Temperature in the cuvette before and immediately after measurements was monitored with a thermo-coupling device.

### RNA degradation measurements

Cy3 labelled RNA oligonucleotide was purchased from Dharmacon (Lafayette, Colorado). RNA degradation by fs laser was measured for 0.15 μM of 38 nt Cy3 labelled RNA substrate (V = 600 μL, 60 mM KCl, 6 mM HEPES-pH 7.5, 0.2 mM MgCl_2_, 5’-GCU UUA CGG UGC UUA AAA CAA AAC AAA ACA AAA CAA AA-Cy3-3’), irradiated with the fs laser (2.6 mW) as described above for 0, 100, 200, 300 and 680 s. RNA degradation by steady-state UV irradiation was measured for 0.15 μM of the 38 nt Cy3 labelled RNA substrate (V = 50 μL,60 mM KCl, 6 mM HEPES-pH 7.5, 0.2 mM MgCl_2_) irradiated in a Stratalinker (Fisher Scientific, 200 mJ/cm^2^) for same time points. Following irradiation, samples were subjected to denaturing PAGE (4-12% Novex NuPage Bis-Tris (Invitrogen), 60 min, 100 V). Samples on the gels were quantified using a Phosphorimager (GE) in fluorescence detection mode. Intact and degraded RNA bands were quantified using the ImageQuantTL (GE) software. The fraction degraded RNA (Frac D) at each time point was calculated according to:

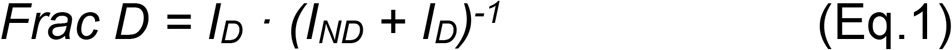

(I_D_: fluorescence intensity degraded RNA, I_ND_: fluorescence intensity non-degraded RNA) Photons absorbed over time (**Extended Data Fig.1b**) were calculated according to ^11,13^

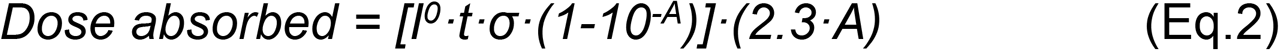

(I^0^ = intensity of incident light in photons cm^−2^ s^−1^; t = duration of irradiation; A = absorbance of protein-RNA solution in Absorbance Units (AU), σ = mean cross section of absorption of nucleic acids). For the fs laser: I^0^ = 2·10^27^ photons cm^−2^ s^−1^ (refs. ^9,13^), A_270_ = 0.99 AU (Absorbance Units of protein-RNA solution), σ = 2.7 × 10^−17^ cm^2^ molecule^−1^ (ref.^13^). For the steady-state UV irradiation (Stratalinker, 400 mJ /cm^2^) I^0^ = 2.10^15^ photons cm^−2^ s^−1^, A_270_ = 0.99 AU, σ = 2.7 × 10^−17^ cm^2^ molecule^−1^ (ref.^13^).

### Protein expression and purification

*Mus musculus* Dazl(RRM) (amino acids 32 – 117) was codon-optimized (Dapcel, OH) for expression in *E.coli*. (**Supplementary Material Table S1**). The DNA construct was chemically synthesized (Genscript, NJ) and cloned into a pET-22b vector with an N-terminal His_6_ – Sumo cleavable tag. Protein was expressed in *E.coli* (BL21) cells overnight at 19°C and purified through Ni^2+^ affinity column ^16^. Samples were dialyzed (20 mM HEPES, pH7.5, 100 mM NaCl), the His_6_-Sumo tag was removed with Sumo protease (Ulp1) at 4°C overnight. Dazl(RRM) protein was further purified by gel filtration chromatography (Superdex 75) equilibrated in 20 mM HEPES (pH 7.5), 100 mM NaCl, 5% (v/v) glycerol. Peak fractions were pooled and concentrated with Amicon ultra centrifugal filters. RbFox(RRM) (amino acids 109-208) and RbFox^mut^(RRM) (amino acids 109-208, R118D, E147R, N151S, E152T mutations) proteins were prepared as described ^15^. Protein concentrations were determined by UV absorbance at 280 nm and validated with Bradford assays.

### RNA-protein affinity measurements by fluorescence polarization

Purified proteins RbFox(RRM), RbFox^mut^(RRM), Dazl(RRM) at different concentrations and corresponding cognate 3’-Cy3 RNAs (20 nM, RbFox: 5′-UCC<u>UGCAUG</u>UUUA-Cy3-3’, Dazl: 5′-UU<u>GUU</u>CUUU-Cy3-3’, cognate motifs underlined; modified RNAs purchased from Dharmacon, Lafayette, Colorado) were incubated for 10 min (20 mM HEPES (pH 7.5), 100 mM NaCl and 0.01% (v/v) NP-40). Solutions were transferred to a 96-well plate (Greiner Bio-one), and fluorescence polarization was measured in a Tecan M1000-Pro microplate reader (Tecan, Switzerland). Plots of the fraction bound RNA vs. protein concentrations were fitted against the quadratic binding equation using KaleidaGraph (Synergy, PA) ^16^.

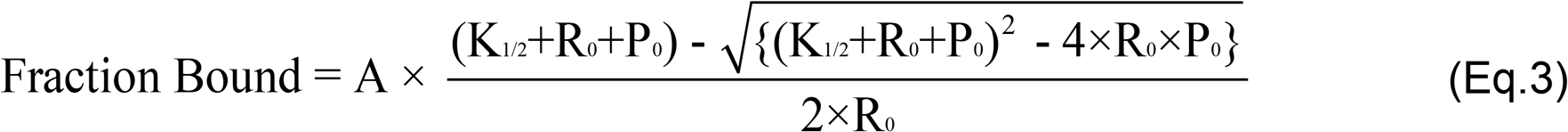

(A: reaction amplitude, K_1/2_: apparent dissociation constant, R_0_: RNA concentration, P_0_: protein concentration)

### fs laser RNA-protein crosslinking *in vitro*

Cy3 labelled RNA oligonucleotides corresponding to cognate sequences for RbFox(RRM) and Dazl(RRM) (described above, 5 nM, final concentration) and protein (10 nM, 50 nM, final concentration) were combined in a cuvette (V = 600 μL, 20 mM HEPES (pH 7.5), 100 mM NaCl, 5% (v/v) glycerol, 25°C) and incubated for 5 min. Longer incubation times did not change results, indicating that equilibrium was reached. The solution in the cuvette was constantly stirred during the reaction (200 rpm), using a magnetic stirbar. Laser power during the measurement was monitored with a photodiode, as described above. Temperature in the cuvette was measured before and after reactions. The RNA-protein mix was irradiated with the UV laser at two different powers (1.0 mW and 2.6 mW, 270 nm). Each timepoint was measured in a separate reaction, avoiding volume changes during the crosslinking experiment. Following crosslinking, samples were removed from the cuvette and stored on ice. Crosslinked and non-crosslinked RNA were separated on denaturing PAGE (4-12% Novex NuPage Bis-Tris gel, 200 V, 45 min). Fluorescence of crosslinked and non-crosslinked RNA in the gels was measured with a Phosphorimager (GE) and quantified with the ImageQuant TL Software (GE). The fraction cross-linked RNA (Frac XL) at each time point was calculated according to:

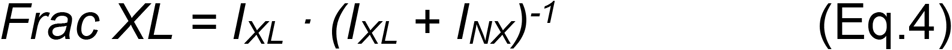

(I_XL_: fluorescence intensity crosslinked material, I_NX_: fluorescence intensity non-crosslinked material). Timecourses at the different protein concentrations and laser powers were globally fit to a two-step kinetic model (**Fig.1a**) using KinTek Global Kinetic Explorer (Kintek, Austin TX). Quality of the global fit was assessed by Chi-squared (Χ^2^) values (**Supplementary Material Fig.S1**). Reported error represents the 95% confidence interval.

### Cell culture

GC-1*spg* cells with inducible DAZL expression were maintained in DMEM high glucose medium (ThermoFisher) supplemented with 10% (v/v) Tet-system approved FBS (Clontech), 100 U/mL penicillin, 100 mg/mL streptomycin, 5 mg/mL blasticidin, and 300 mg/mL Zeocin (all from ThermoFisher) at 37°C, 5% (v/v) CO_2_ (ref.^17^). Doxycycline induction of Dazl was performed and lysates for generation of cDNA libraries and quantification of Dazl levels were prepared as described ^17^. Equal amounts of protein were run on a SDS-PAGE (10% NEXT Gel, Amresco) and transfered to a PVDF membrane. Western blotting was performed with anti-Dazl (Rabbit; 1:5000, US Biological) and anti-Hsp90 (Rabbit;1:10,000; US Biological) antibodies. Chemiluminescence was quantified with the ImagequantTL software.

### fs laser crosslinking of GC-1 cells

GC-1*spg* cells (with doxycyline induction of Dazl expression) were grown in 150 mm plates to 70% confluency. Cells were rinsed with 2 mL PBS (per plate), scraped, re-suspended in 600 μL PBS, transferred to the quartz cuvette and stirred with a magnetic stir bar (described above). Crosslinking of the cell suspension was performed as described above at two laser powers (1.0 mW, 2.6 mW) in separate experiments for 30, 180 and 680 s (25°C). Each crosslinking reaction contained a constant number of cells (6·10^5^). To generate sufficient material for timepoints with low crosslinking yield, multiple identical experiments were conducted and pooled. Temperature in the cuvette was measured before and after crosslinking (increase was less than 1°C after 680 s). Cell integrity after crosslinking was measured by trypan-blue staining ^39^ and cell counting in a hemocytometer. After crosslinking, cell suspensions were pelleted at 1,000 g for 5 min (4°C). The pellet was suspended in PBS (3x dry volume). Cells were pelleted again (1,000 g for 5 min), the supernatant was removed, and pellets were frozen and stored at −80°C until further processing.

### cDNA library preparation

Cell lysates for each sample were split into two aliquots (A1, A2). RQ1 DNase (PromegaM6101) and RNAse A (USB70194Y) were added at 1:100 (A1) and 1: 20,000 (A2). Over-digested sample (A1) confirmed the size of the Dazl-RNA radioactive band on SDS-PAGE gel. The under-digested cell supernatant from the under-digested sample (A2, equivalent to ~150 mg of cell lysate) was mixed with protein G Dynabeads (ThermoFisher 10009D) with anti Dazl antibody (Rabbit; 1:5000) in separate Eppendorf tubes for each sample (N = 16). Samples were treated with CIP (Roche712023). RNA linker ligation and PNK (NEBM0201S) treatment were performed as described ^17^. The supernatants were loaded onto separate Novex NuPAGE 4-12% Bis-Tris gels, and crosslinked material was transferred to a nitrocellulose membrane. Samples were located on the membrane by autoradiography and RNA-Dazl complexes at 50 – 70 kDa (Dazl molecular weight; 37 kD) were cut. Nitrocellulose fragments were treated with proteinase K (Roche1373196) Dazl bound RNA was isolated, reverse transcribed (SuperScript III (Invitrogen18080051), circularized and amplified to obtain 16 cDNA libraries. The RT primers used contained iSP18 spacers and phosphorylated 5’ end for circularization of first strand cDNA to generate PCR template without linearization ^17^. Unique molecular identifiers (UMIs, randomized barcodes, 11 nt with 4 nt random nucleotides) were used to determine PCR amplification artifacts (primer sequences: **Supplementary Material Table S2**). cDNA diversity in each library was tested before next generation sequencing by cloning cDNA from each library into pBS plasmid, subsequent transformation in competent cells, colony PCR and DNA sequencing. Illumina Sequencing for all cDNA libraries was performed at the Case Western sequencing core facility.

### Measurement of bulk crosslinking

For each KIN-CLIP library, cells were cross linked and cell lysate was prepared as described above. 200 μL aliquots (equivalent to 150 mg of cell lysate) for each KIN-CLIP sample were treated with RQ1 DNase and RNAse (at 1: 20,000) as described above. Treated lysates were centrifuged in a pre-chilled ultra-centrifuge, polycarbonate tubes, TLA 120.2 rotor at 30,000 rpm, 20 min, 75 μl of the supernatant were removed and RNA was 5’-radiolabeled with PNK. Samples were run on a SDS-PAGE gel and transferred to a nitrocellulose membrane. The radioactivity was measured by quantifying the intensity of the radioactive bands (using ImageJ software). Lane background was used to normalize the band intensities.

### KIN-CLIP read processing, refinement and mapping

Raw sequencing reads were assessed for quality (FastQC, https://www.bioinformatics.babraham.ac.uk) and de-multiplexed. Low-quality reads were removed if ≤ 80% of sequenced bases in a read had a PHRED quality score of ≤ 25. De-multiplexing and read filtering was performed with the FASTX-Toolkit (http://hannonlab.cshl.edu/fastx_toolkit/) using standard commands ^40^. Filtered reads were stored in FASTQ format. Barcode and UMI (randomized 4nt sequence) were kept appended to line 1 of the FASTQ for each read.

Read duplicates, as identified by UMIs were collapsed into a single read. Linkers and concatamers were removed with the FASTX-Toolkit (http://hannonlab.cshl.edu/fastx_toolkit/), using permutations (N = 25) of linker sequences as target. Reads with ≥ 15 nt were retained for subsequent analysis. Processed reads were aligned against the mouse genome (mm10) by using bowtie2 ^41^ with the following settings for a 50 bp sequencing run: Number of mismatches allowed in seed alignment during multi-seed alignment = *1*, length of the seed substrings to align during multi-seed alignment = *15*, set a function governing the interval between seed substrings to use during multi-seed alignment = *S,1,0.50,* function governing the maximum number of ambiguous characters (N’s and/or ‘.’s) allowed in a read as a function of read length = *L,0,0.15*, disallow gaps within this many positions of the beginning or end of the read = *4*, set a function governing the minimum alignment score needed for an alignment to be considered ‘valid’ = *L, −0.6, −0.6*, set the maximum (‘MX’) and minimum (‘MN’) mismatch penalties, both integers = *6,2,* sets penalty for positions where the read, reference, or both, contain an ambiguous character such as *‘N’ = 1,* gap opening penalty = *5*, gap extension penalty = *3*, attempt that many consecutive seed extension attempts to ‘fail’ before Bowtie 2 moves on, using the alignments found so far = *20*, set the maximum number of times Bowtie 2 will ‘re-seed’ reads with repetitive seeds = *3*. End-to-end alignment mode was used. Only uniquely mapped reads were retained. To evaluate the stringency of filtering and sequence alignment, the fraction of uniquely mapped tags over all mapped reads was assessed ^40^ by employing different permutations of read mapping parameters described above. In total, 55 parameter permutations for mapping were tested. The setting yielding the largest number of uniquely mapped reads is shown above. The *BAM* index of mapped reads corresponding to the 16 KIN-CLIP libraries was then converted to *BED*/*bedgraph* using the standard command line version of –bedtools (V2.29.1) and –samtools (V1.10) ^42^. Bedgraph files were visualized in the IGV ^43^.

### Identification of KIN-CLIP peaks

Genomic coordinates of the 5’-terminal nucleotide (5’nt) of every mapped read were obtained. Adjacent 5’nt were summed at single nucleotide resolution level by creating a sliding window of 11nt (stride = 1, steps = 5nt on either side or until no new reads were detected), with the 5’nt position at the center. Crosslinking peaks were defined by plotting the distribution of the count of 5’nt reads in these windows for every location. The peak apex represents the coordinate for the crosslinking peak and the associated coverage value. Error ranges for coverage values corresponding to each crosslinking peak were defined as the 95% confidence interval from the apex of crosslinking peaks. Coordinates of crosslinking peaks present in all KIN-CLIP libraries, except at the zero timepoint were used to define Dazl binding sites for further analysis. For peaks with coverage at the zero timepoint (~0.2% of peaks), the peak value at *t* = 0 was subtracted from the KIN-CLIP peaks. Coverage values for each Dazl binding site were converted into a concentration equivalent by normalizing to the amount of bulk crosslinked RNA for each KIN-CLIP library (**Supplementary Material Table S6**). The normalized read coverage values were used for calculating kinetic parameters and other subsequent analyses.

### Analysis of read distribution

To annotate KIN-CLIP Dazl binding sites, RefSeq coding regions, 5’UTRs, 3’UTRs, ORF, introns, and RNA types were obtained from the UCSC genome browser and intersected individually with KIN-CLIP binding site coordinates using *bedtools*.

### CITS analysis and sequence enrichment

Crosslink Induced Truncation Site (CITS) analysis was performed as described ^28,29^. Enrichment of motifs at and around CLIP regions was performed using the EMBOSS tool Compseq ^44^, R package ‘randomizeR’ ^45^ and ‘Random’ ^46^ module in Python. To generate z-scores, shuffled control sets were generated for each dataset analyzed using Random module available in Python (Shuffle N = 10,000).

### Distribution of Dazl-RNA contacts in 3’UTRs

Metagene analysis of Dazl-3’UTR interactions was performed on 3’UTRs as defined by PolyA-Seq ^17^). To define 3’UTR length, coordinates from Refseq and Ensembl ^30,31^ were matched with PolyA-Seq data ^17^. For transcripts with multiple 3’UTR length annotations, coordinates for the longest 3’UTR were utilized. 3’UTRs that overlapped with intron sequences annotated in either RefSeq or Ensembl were omitted. To calculate distances of binding sites to PAS and stop codons, the distance between coordinates for each KIN-CLIP binding relative to the Stop codon and to the PAS (10 nt window) was measured. For each 3’UTR, the random distribution of binding sites was determined by scrambling all Dazl binding sites (1,000 times) in that 3’UTR into all probable 10 nt bins in that 3’UTR and obtaining the average.

### Calculation of kinetic parameters

Kinetic parameters were calculated from normalized peak coverage values for each Dazl binding site (N = 10,341). A Dazl binding site was defined by the presence of more than 7 normalized sequencing reads in the library for the (4.2xDazl, 2.6 mW laser) 680 s timepoint, within 11 nucleotides of the peak apex for the binding site in all libraries. Kinetic parameters were calculated according to two different approaches: (i) a numerical and (ii) an analytical method. Parameters from both methods were averaged for subsequent data analysis (**Extended Data Fig. 3**).

### Numerical approach

The numerical approach to calculate kinetic parameters is based on numerically fitting crosslinking timecourses to the differential equations describing the Dazl-RNA binding and crosslinking process (**Fig.1a**), according to:

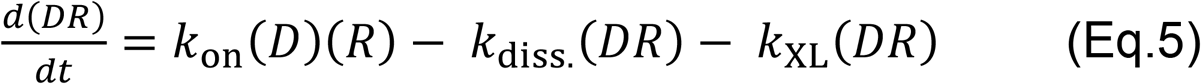

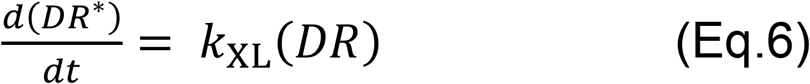

(DR: concentration of non-crosslinked Dazl-RNA complex (for each binding site), DR*: concentration of crosslinked Dazl-RNA complex (for each binding site), D: Dazl concentration, R: RNA concentration (binding site), *k*_on_: association rate constant, *k*_diss._: dissociation rate constant, *k*_XL_: crosslinking rate constant).

Because concentrations of free Dazl and RNA in the cell are experimentally inaccessible, the second order association process (*k*_on_) was treated as pseudo-first order reaction at each of the two Dazl concentrations. Accordingly, we calculated a pseudo first order rate constant for each Dazl concentration (*k*_on_^(1xDazl)^, *k*_on_^(4.2xDazl)^), and *k*_diss._, *k*_XL_ ^(1mW)^ and *k*_XL_ ^(2.6mW)^ for each binding site. Numerical fitting of timecourses of normalized read coverage for each binding site (**Fig.2c**) was performed in R with packages deSolve (with ODE function) ^47^, ggplot2 ^48^, reshape2 ^49^ and rmarkdown ^50^.

The fitting strategy encompassed two steps: (i) estimation of parameter ranges following a sequential parameter estimation procedure ^51^ and (ii) fitting the timecourses using estimated parameter ranges as input (**Supplementary Material, Scheme 1**). Estimation of parameter ranges was also performed in two steps, (i,a) *initial* parameter range estimation for *k*_on_^(1xDazl)^, *k*_on_^(4.2xDazl)^, *k*_diss._, *k*_XL_^(1mW)^ and *k*_XL_^(2.6mW)^, and (i,b) refinement of *initial* parameter range estimates to obtain *final* parameter range estimates (**Supplementary Material, Scheme 1**). To estimate *initial* parameter ranges, timecourses from reactions with 4.2xDazl at high laser power (2.6 mW) and low laser power (1mW) were fit separately. Starting values were based on the kinetic parameters measured *in vitro* (**Fig.1**; *k*_on_^(1xDazl)^ = 0.0001 s^−1^, *k*_on_^(4.2xDazl)^ = 0.0001 s^−1^, *k*_*diss.*_ = 1 s^−1^, *k*_XL_ ^(1mW)^ = 10 s^−1^ and *k*_XL_ ^(2.6mW)^ = 1 s^−1^. Use of significantly different starting values did not yield acceptable fits for the majority of binding sites). This step provided average initial values for *k*_on_^(4.2xDazl)^ and *k*_diss._ as well as initial values for *k*_XL_^(1mW)^ and *k*_XL_^(2.6mW)^. Next, timecourses from reactions at 1xDazl at high laser power (2.6mW) and low laser power (1mW) were fit separately, yielding average initial values for *k*_on_^(1xDazl)^ and *k_diss._* and initial values for *k*_XL_^(1mW)^ and *k*_XL_^(2.6mW)^. *k*_XL_^(1mW)^ and *k*_XL_^(2.6mW)^ were then averaged. This process was performed for each binding site until the Χ^2^ was minimized (no change in Χ^2^ for 4 consecutive cycles) or 1,000 fitting cycles were completed. The process provided 10,341 × 5 parameter values, which were plotted as distribution (10,341 values for each parameter). The *initial* parameter range estimate represents the 95% confidence interval from the mean of the distribution for *k*_on_^(1xDazl)^, *k*_on_^(4.2xDazl)^, *k*_diss._, *k*_XL_^(1mW)^ and *k*_XL_^(2.6mW)^.

To refine parameter range estimates to obtain *final* parameter range estimates, the initial parameter range estimates were used as input to fit multiple, random subsets of 2,000 randomly selected binding sites. 10,000 iterations, each with a unique random subset of 2,000 binding were performed. Each iteration yielded a distribution. All 10,000 distributions were superimposed and the median apex of all distributions was identified. The *final* parameter range estimates represent the 95% confidence interval from the median apex of the averaged distributions. The *final* parameter range estimate was about 35% smaller than the *initial* parameter range estimate.

The estimated parameter ranges were used as input for fitting of the timecourses (**Supplementary Material, Scheme 1**). We fitted timecourses for reactions at 4.2xDazl at the different laser powers (1 mW, 2.6 mW), varying linked *k*_on_^(4.2xDazl)^ and *k*_diss._ (which do not scale with laser power), and differing *k*_XL_^(1 mW)^ and *k*_XL_^(2.6 mW)^. We then fit timecourses at 1xDazl at both laser powers, varying linked *k*_on_^(1xDazl)^ and *k*_diss._, and differing *k*_XL_^(1 mW)^ and *k*_XL_^(2.6 mW)^. Utilizing parameters obtained from these two steps we fit all 4 timecourses linking *k*_on_^(4.2xDazl)^ and *k*_on_^(1xDazl)^ for differing laser powers, linking *k*_XL_^(2.6 mW)^, *k*_XL_^(1 mW)^ for differing Dazl concentrations and linking *k*_diss._ for all conditions. The process of fitting all 4 timecourses for each binding site was repeated 642 times, after which χ^2^ did not show significant fluctuation (< 5% for 4 consecutive cycles). Obtained rate constants were used as final kinetic parameters for the numerical approach (**Extended Data Fig.3b-d**).

Fitting quality was assessed by calculating chi-squared (χ^2^) for each binding site, the overall cumulative reduced chi-squared (χv^2^) and the coefficient of determination/R^2^ (COD) according to:

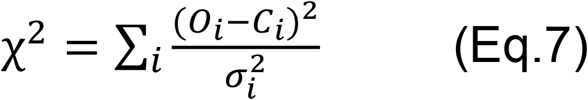

(*O*: observed value, *C*: calculated value for each binding site (i). 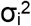 is the squared variance between data points *O, C*);

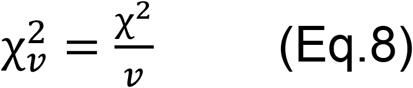

[*v*: degree of free; equals *(n − m)*, with *n*: number of observations (*n = 16*), *m*: number of fitted parameters (*m = 5*)].

The coefficient of determination/R^2^ (COD) was calculated using the standard method as described ^52^. The COD describes correlation between calculated and observed timecourses. For the last fitting cycle, COD = 0.92, *Χ*_*v*_^*2*^ = 0.043 (**Extended Data Fig.3c**).

### Analytical approach

The analytical approach to calculate kinetic parameters is based on fitting of crosslinking timecourses to explicit solutions of the system of differential equations (Eqs.5,6) for the kinetic scheme (**Fig.1a)**. To solve the system of differential equations, we considered that at any given time *(t)* during crosslinking, the accessible fraction of a given Dazl binding site is either free *(R)*, occupied *(DR)* or crosslinked *(DR*)*:

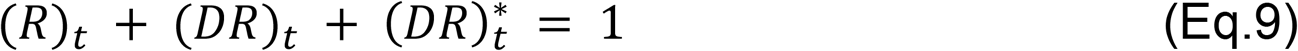

In addition, at *t* → *∞*, 100% of the accessible fraction of a given Dazl binding site is crosslinked. As described for the numerical approach, the second order association process (*k*_on_) was treated as pseudo-first order process at each Dazl concentration.

Treating second order association process (*k_on_*) as pseudo-first order process, considering Eq.9 and rearranging Eq.5 yields:

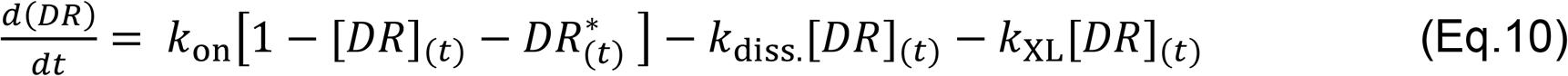

Before crosslinking (t = 0), at steady-state of the binding reaction,

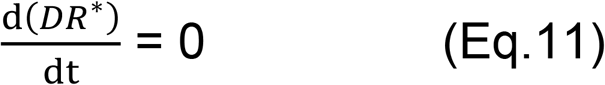

because

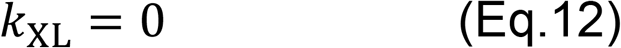

From Eq.5, we thus obtain:

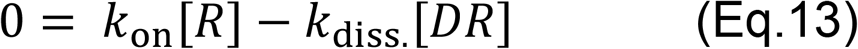

which yields, after rearranging,

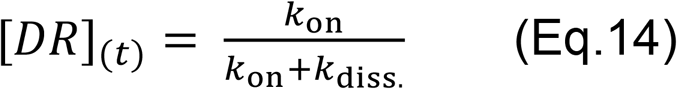

At t = ∞, crosslinking is complete, and thus

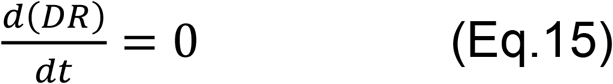

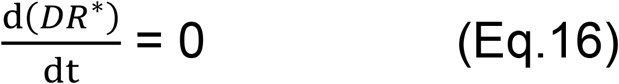

The boundary limits are:

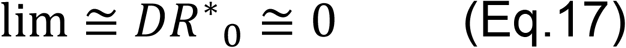

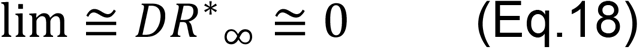

Equations 11-18 define the boundary conditions.

Crosslinking timecourses represent amount of crosslinked material at a given time (t), expressed as normalized coverage value for each binding site [DR*]_(t)_. [DR*]_(t)_ depends on amount of Dazl-RNA complex [DR] at the time (t) (Eq.6) and thus on *k*_on_, *k*_diss._ and *k*_XL_. Absolute concentrations of [D], [R] and [DR] are not known in our system. To extract *k*_on_, *k*_diss._ and *k*_XL_ for each binding site from the crosslinking timecourses we integrate Eq.6 after appropriate substitution of [DR]. To accomplish this, we take a second differential of Eq.10, considering the boundary conditions (Eq.11-18). We obtain the general solution of the second order differential equation:

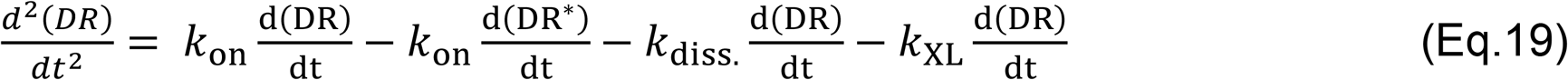

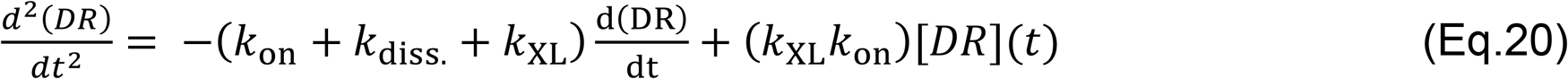

Equation 20 is a constant coefficient, homogenous, linear, second order differential equation with two independent solutions (*y1*, *y2*) ^53^:

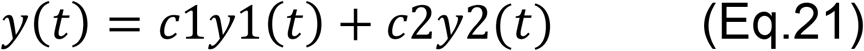

The coefficients *c1* and *c2* (by the principle of superposition) ^54^ are obtained after providing the boundary conditions from equations 11-18. We identify a function *y* where a constant multiplied by its second derivative *y’’* plus another constant times *y’* plus a third constant multiplied by *y* equals zero ^54^.

The exponential function

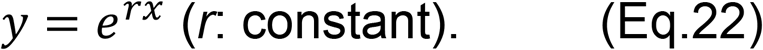

has the property that its derivative is a constant multiple of itself:

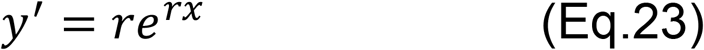

Furthermore,

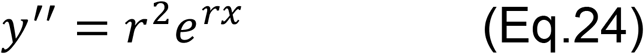

Substituting these expressions into (Eq.20), we obtain:

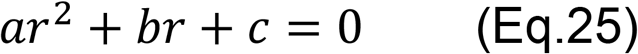

Equation 25 is the auxiliary (characteristic) equation of the differential equation 20 (ref. ^55^). The equation is transformed into an algebraic equation by replacing

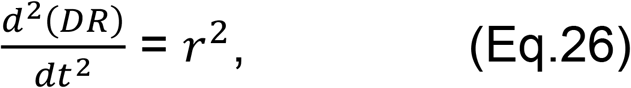

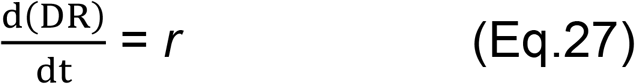

and [*DR*] by 1.

The roots of Eq.25 are found by factoring ^55^:

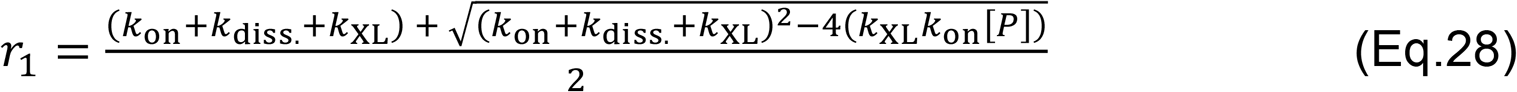

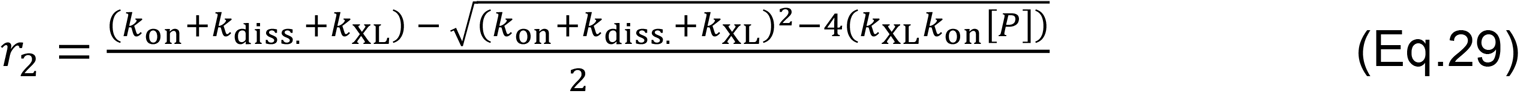

With Eq.21-29, the general solution of Eq.20 is ^56^:

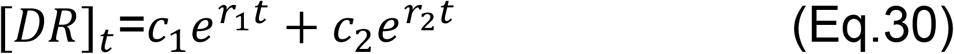

To obtain our observable *[DR*]_(t)_*, we integrate Eq.6 under consideration of the boundary conditions (Eqs.11-18):

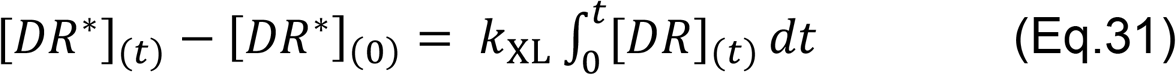

Substituting *[DR]_t_* from Eq.30 yields

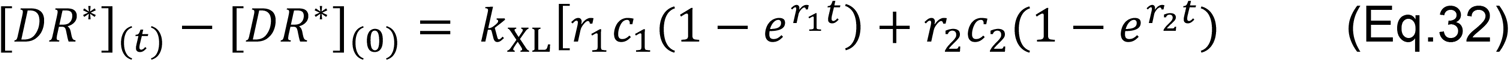

Substituting *c1* and *c2* by providing the boundary conditions (Eqs.11-18) and considering (Eqs.21-29), we obtain:

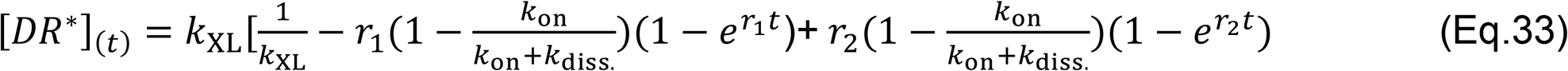

Equation 33 is an explicit nonlinear equation of the form:

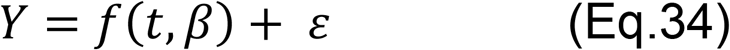

*t* = *(t_1_*, *t_2_*, *t_n_)* are the independent variables (the normalized read coverage values at different timepoints), *β* = *(β_1_*, *β_2_*, … …. . *β_n_)’* are the parameters (*k*_on_ = *k*_on_^4.2x #i^, *k*_on_^1x #i^*, k*_XL_ = *k*_XL_^2.6mW #i^*, k*_XL_^1mW #*i*^ and *k*_diss._ = *k*_diss._^#i^, where *#i* represents the crosslinking conditions. *ε* is the fitting error between observed and expected timecourses. *f(t, β)* represents the functional relationship between t, β and Y.

Equation 33, adapted to the different Dazl concentrations and different laser powers was used to fit the crosslinking timecourses for each binding site. The resulting equations represent the non-linear model:

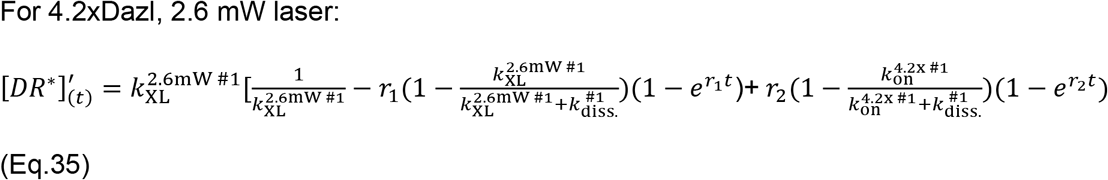

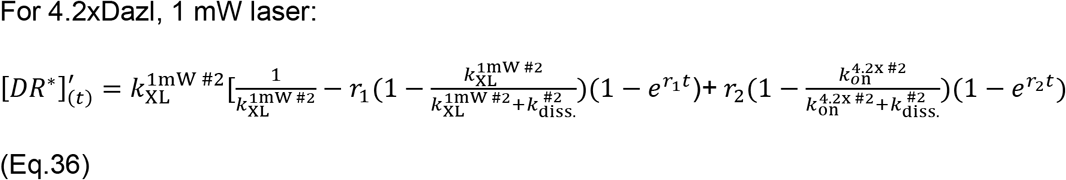

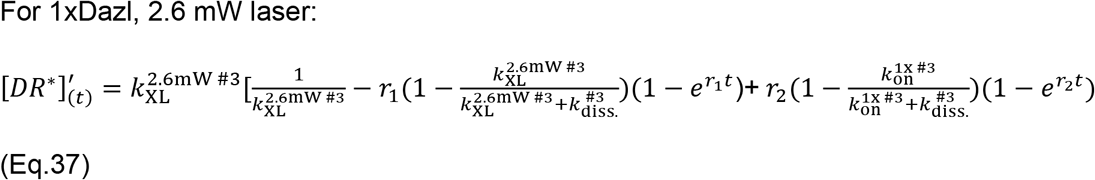

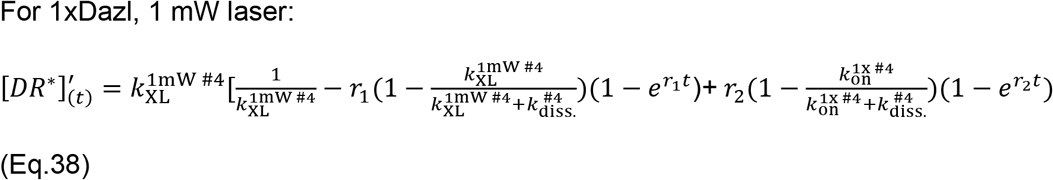

r1 and r2 are:

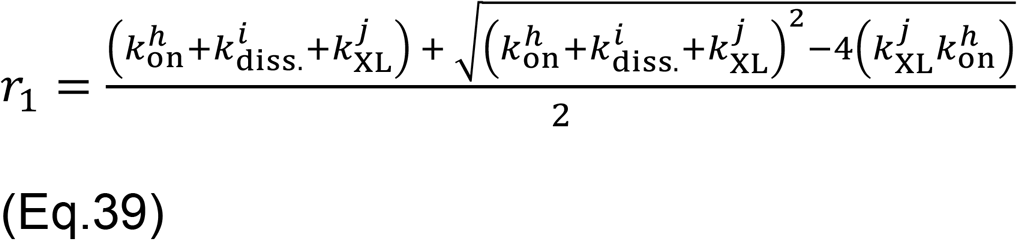

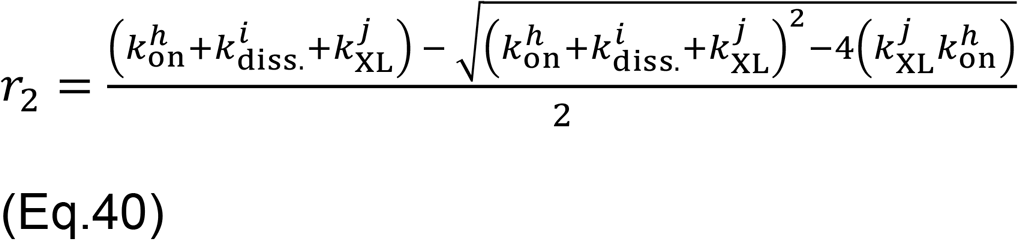

*h* represents 4.2xDazl #1 (Eq. 35), 4.2xDazl #2 (Eq.36), 1xDazl #3 (Eq.37) and 1xDazl #4 (Eq.38). *i* represents #1 (Eq.35), #2 (Eq.36), #3 (Eq.37) and #4 (Eq.38). *j* represents 2.6 mW #1 (Eq.35), 1 mW #2 (Eq.36), 2.6 mW #3 (Eq.37) and 1 mW #4 (Eq.38).

Timecourses for 4.2xDazl at high laser (2.6 mW), 4.2xDazl at low laser (1mW), 1xDazl at high laser power (2.6 mW) and 1xDazl at low laser power (1mW) were separately fit to the non-linear model (**Supplementary Material Scheme 2**).

A matrix of initial parameters was obtained,

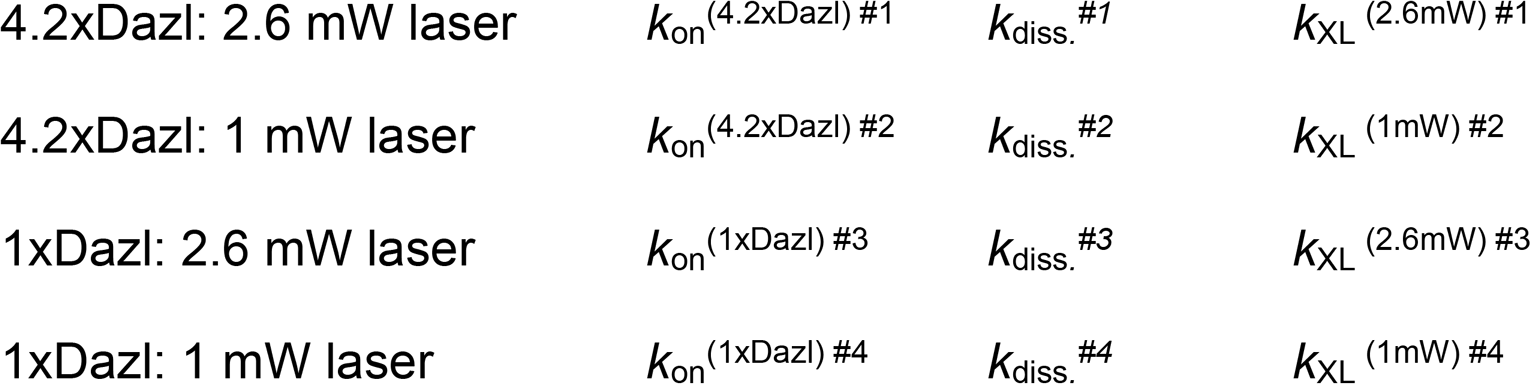

Next, a global datafit for all four timecourses (#1-4) for an individual binding site was performed. Initial parameters were iteratively adjusted, considering the following criteria:

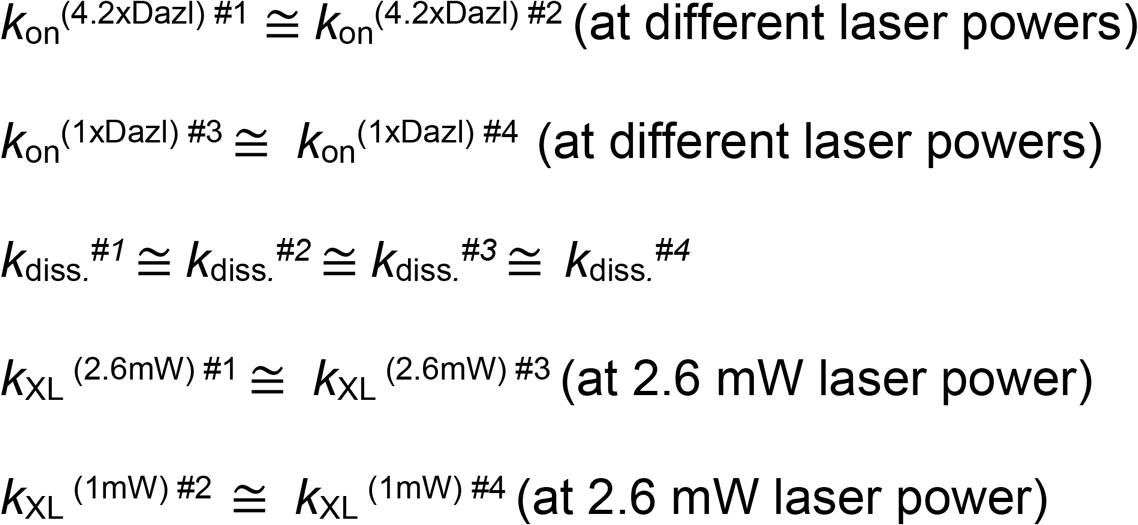

Fits were repeated until the best fit was reached (no change in Χ^2^ for 4 successive fittings), as measured by Chi-squared Χ^2^ minimization, according to:

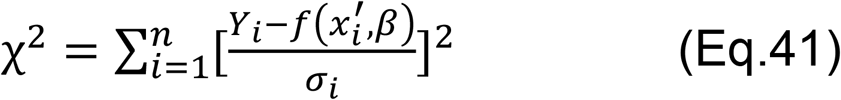

*x_i_’* is the row vector for the *ith* (*i* = 1, 2, …, n; n = 10,341) observation. *β* is the parameter under consideration. *Y_i_* is the estimated parameter value for the *i_th_* (i = 1, 2, …, n; n = 10,341) observation. *σ_i_* is the variance between observed and estimated parameter values. *f(x_i_’, β)* represents the function for which *x_i_’* and *β* are measured.

Obtained parameters were further refined by additional rounds of fitting using the analytical, Levenberg-Marquardt (L-M) least squares algorithm, which combines the Gauss-Newton and the steepest descent method ^57^. Utilizing the values obtained above, parameters for timecourses at 4.2xDazl at high laser power (2.6 mW) and low laser power (1mW) were adjusted together. *k*_on_^(4.2xDazl) #2^ was increased or decreased (depending on initial values for a given binding site) in small increments (∂b) in order to move *k*_on_^(4.2xDazl) #2^ closer to *k*_on_^(4.2xDazl) #1^. ∂b was set as 5% of *k*_on_^(4.2xDazl) #2^ for a given binding site. Following each increment, the timecourse was fitted to the non-linear model and Χ^2^ calculated. *k*_diss._^*#2*^ and *k*_XL_ ^(1mW) #2^ were floated during the fitting. If Χ^2^ (b + ∂b) ≥ Χ^2^ (b) for >3 consecutive fitting cycles, *k*_on_^(4.2xDazl) #1^ was increased or decreased (depending on initial values) in small increments to improve fitting. This fitting procedure was repeated for N = 642 cycles.

Next, the parameters for timecourses at 1xDazl at high (2.6 mW) and low laser power (1mW) were adjusted, providing *k*_on_^(4.2xDazl) #1^, *k*_on_^(4.2xDazl) #2^, *k*_on_^(1xDazl) #3^ and *k*_on_^(1xDazl) #4^. Keeping the adjusted *k*_on_ constant (floating *k*_XL_), were subsequently adjusted *k*_diss._^*#1*^, *k*_diss._^*#2*^, *k*_*diss*._^*#3*^ and *k*_*diss*._^*#4*^ (within 25% range of each other). Finally, *k*_XL_ ^(2.6mW) #1^ and *k*_XL_ ^(2.6mW) #3^ were adjusted by increasing or decreasing *k*_on_^(4.2xDazl) #1^ and *k*_on_^(4.2xDazl) #3^ in small increments (∂b ≤5% of parameter values) while maintaining *k*_on_^(4.2xDazl) #1^ > *k*_on_^(4.2xDazl) #3^. Additionally, *k*_diss._^*#1*^ and *k*_diss._^*#3*^ were increased or decreased in increments of ∂b ≤1%. The same process was performed for adjusting *k*_on_^(4.2xDazl) #2^ and *k*_on_^(4.2xDazl) #4^. Every parameter adjustment cycle was repeated 642 times after which Χ^2^ values computed in 4 successive iterations showed fluctuations of less than 5% for > 95% of binding sites.

### Calculation of binding probabilities

The binding probability (P) describes the probability by which the accessible fraction of a given binding site is bound by Dazl. P for each Dazl concentration was calculated according to:

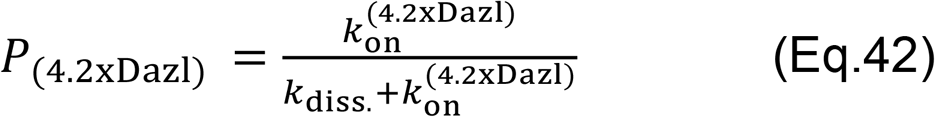

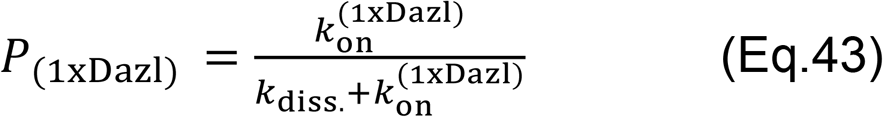

### Calculation of fractional occupancy

The fractional occupancy (Φ^max^) describes the fraction of a given binding site that is occupied by Dazl extrapolated to saturating concentrations. Φ^max^ is a measure of binding site accessibility during the course of the experiment. Φ^max^ = 1 indicates complete accessibility, decreasing values indicate decreasing accessibility. Φ^max^ was calculated by plotting the maximal amplitude (α^max^: probability of Dazl bound to the fraction of a given binding site that is accessible during the course of the experiment, extrapolated to saturating concentrations of Dazl) vs. level of the corresponding transcript (L, in RPKM) (**Supplementary Material Figure S2**). Φ^max^ corresponds to the slope of the plots, and was calculated according to:

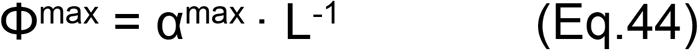

Reported Φ^max^ values were normalized to a scale of zero to 1. To define α^max^, apparent association rate constants at both Dazl concentrations *k*_on_^(4.2xDazl)^, *k*_on_^(1xDazl)^ were plotted against the relative cellular Dazl concentrations ([Dazl]^rel^, **Supplementary Material Figure S2**).

For binding sites where *k*_on_^(4.2xDazl)^, *k*_on_^(1xDazl)^ increased linearly with [Dazl]^rel^:

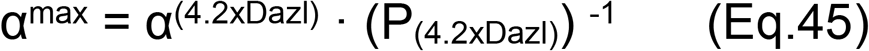

(α^(4.2xDazl)^: normalized read density at the 30s time point for the timecourse with 4.2xDazl and 2.6 mW laser power for a given binding site, P_(4.2xDazl)_: binding probability at 4.2xDazl (Eq.42). For binding sites where *k*_on_^(4.2xDazl)^, *k*_on_^(1xDazl)^ increased with [Dazl]^rel^ in a hyperbolic fashion, we determined the maximal apparent binding rate constant *k*_on_^max^ by fitting the plot of *k*_on_^(4.2xDazl)^, *k*_on_^(1xDazl)^ vs. [Dazl]^rel^ to:

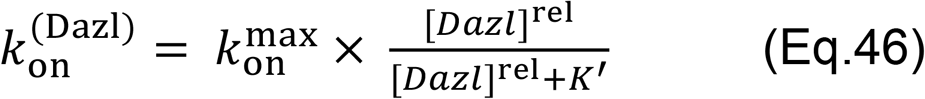

(*k*_on_^(Dazl)^: *k*_on_^(1xDazl)^, *k*_on_^(4.2xDazl)^, K’: apparent relative binding constant)

The binding probability extrapolated to [Dazl]^rel^ saturation (P_max_) is:

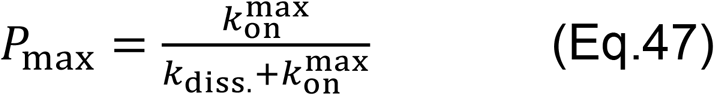

and

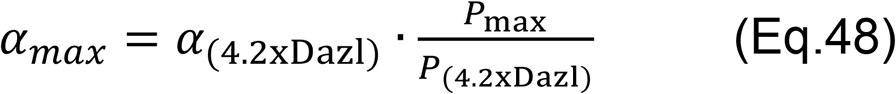

A plot was defined as hyperbolic if *k*_on_^max^ < 4·*k*_on_^(4.2xDazl)^.

### Analysis of Variance (ANOVA)

One-way ANOVA was calculated in R using libraries – car ^58^. Mean square differences between and within groups were calculated. Obtained F values were compared with the critical value in the F table to obtain p values ^58^. Inter-group differences were significant (p < 0.05) when the F value exceeded the critical F value for the given degrees of freedom ^59^.

### Determination of distances between neighboring binding sites

Distances between neighboring binding sites (genomic coordinates: mm10) were calculated between first and last read coordinates of adjacent peaks recorded with a sliding window, (start: l = 0 (chr1), length = 2 nt, stride = 1 nt) for each transcript. The number of inter-site distances for a given value was divided by the overall number of distances to yield the normalized frequency (**Fig.3a**). The random distribution of inter-site distances was obtained by Monte Carlo simulations (**Fig.3a**). A random binding site was defined as a genomic coordinate encompassing a non-overlapping 5 nt long sequence (in the entire mouse transcriptome, **Fig.3a**) within 500 nt of PAS, or excluding 500 nt proximal to PAS, (**Extended Data Fig.5**). 10,341 binding sites were randomly distributed over these windows, their distribution was recorded and plotted as described above. Monte Carlo simulations (Vignette package in R ^60^ were carried out 1,000 times. Obtained distributions were averaged and plotted (**Fig.3a**).

### Dazl cluster definition and distribution

A cluster of Dazl binding sites was defined by an inter-binding site distance of < 40 nt and absence of additional binding sites < 120 nt around the cluster. The distribution of clusters in 3’UTRs (**Fig.3b**) was calculated by dividing the 3’UTRs in 100 nt bins, starting at the PAS. The number of clusters in each bin was counted and the cumulative frequency of clusters with different numbers of binding sites was plotted against the 3’UTR bins.

### Calculation of cumulative and differential binding probabilities

Cumulative binding probabilities (ΣB) for each cluster of Dazl binding sites were calculated according to:

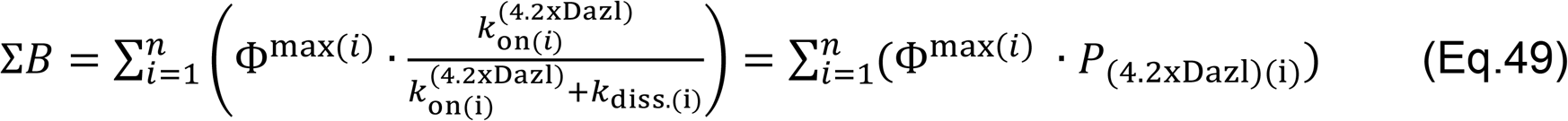

[*n*: number of binding sites in a given cluster; i: individual binding site, Φ^max(i)^: fractional occupancy for the binding site (i); *k*_on(i)_^(4.2xDazl)^: association rate constant at 4.2xDazl for the binding site (i); *k*_diss.(i)_, dissociation rate constant for the binding site (i); P_(4.2xDazl)(i)_: binding probability at 4.2xDazl) for the binding site (i)].

The differential cumulative binding probabilities (ΔΣB) for each cluster of Dazl binding sites were:

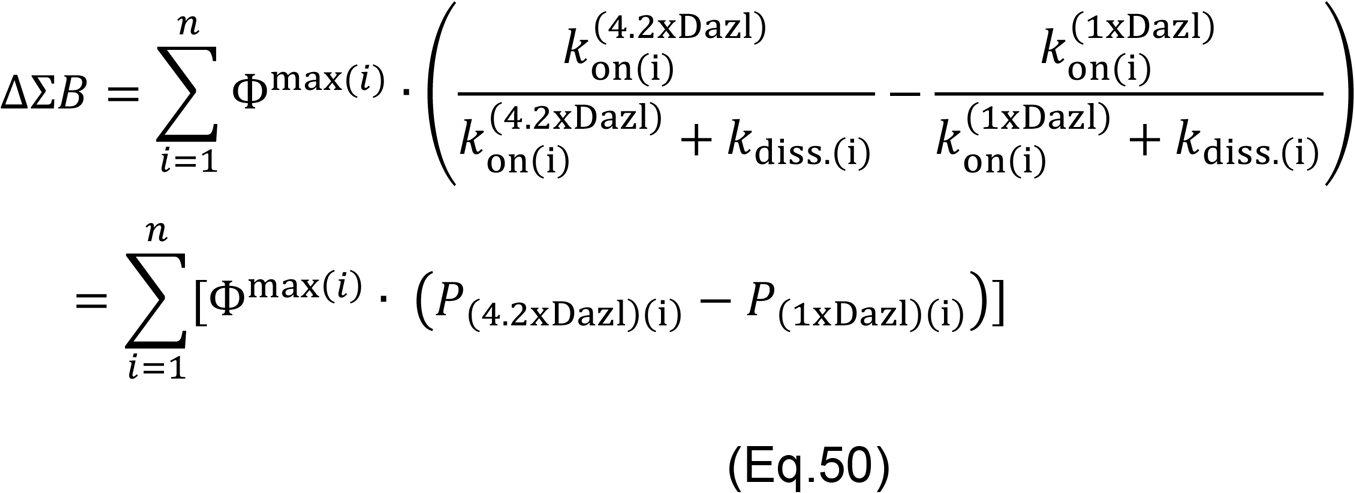

[Variables as above, *k*_on(i)_^(1xDazl)^: association rate constant at 1xDazl for the binding site (i); *k*_diss.(i)_, dissociation rate constant for binding site (i); P_(1xDazl)(i)_: binding probability at 1xDazl for binding site (i)].

### Ribosome Profiling and RNA-seq

Ribosome profiling and RNA–seq, performed in biological triplicates at both Dazl concentrations was described ^17^. Deposited sequencing data (GEO: GSE108997) were analyzed as described ^17^. Averages from the triplicate datasets were used for subsequent data analysis.

### Definition of functional mRNA classes

Changes in ribosome protected fragments (ΔRPF) from 4.2xDazl to 1xDazl (RPKM) and changes in transcript levels (ΔRNA) from 4.2xDazl to 1xDazl (RPKM) for each transcript with a Dazl binding site, represented in all ribosome profiling and RNA-seq datasets were plotted (**Fig.4b**). Low abundance transcripts (RPKM_4.2xDazl_ < 6.0) were removed. ΔRPF and ΔRNA distributions for Dazl bound transcripts were divided into terciles, based on testing the significance (p < 0.05) of the deviation from the mean (H = High; ΔRPF = 1.063, ΔRNA = 1.088, M = Medium; 1.063 ≤ ΔRPF ≤ 0.913, ΔRNA = 1.088 ≤ ΔRPF ≤ 0.974, L = Low; ΔRPF = 0.913, ΔRNA = 0.974). Terciles for ΔRPF and ΔRNA yield nine functional mRNA classes (**Fig.4b**). The HL and LH classes contained too few transcripts (< 10) for meaningful examination and were therefore not considered in subsequent analyses. The MM class was not further considered because neither ribosome occupancy nor transcript level changed significantly upon changes in Dazl concentration.

### Enrichment Analysis

Statistical enrichment of clusters with high, medium and low cumulative binding probabilities (ΣB, **Fig.4a**) in transcripts belonging to each of the functional mRNA classes HH, HM, MH, ML, LM and LL (**Fig.4c**), was calculated with the cumulative distribution function (CDF) of a hypergeometric distribution ^61^ according to:

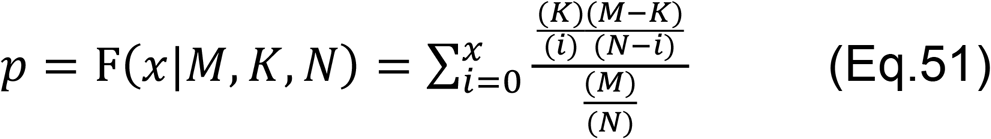

(M: number of total clusters in Dazl bound transcripts, K: number of clusters in each functional mRNA class (HH, HM, MH, LM, LL and ML), N: number of clusters in a given ΣB tercile (H, M, L), i: number of clusters with a ΣB tercile in a given functional mRNA class (for example, number of clusters with high ΣB in HH functional mRNA class). x represents a cluster and F (x|M,K,N) is enrichment of x given M, K and N (by Fishers’ t-test represented as F). p is the hypergeometric p value of enrichment, based on the F-test ^61^) Hypergeometric tests were performed with *Scipy* hypergeom module ^62^ in Python 3.6.5.

### PCA and t-SNE

A data matrix (X) with the seven features of Dazl clusters and of transcripts with Dazl binding sites in 3’UTR (number of clusters in 3’UTR, ΣB, ΔΣB, number of binding sites in a cluster, UTR length, proximity to PAS, transcript level), corresponding to each transcript, was generated. In transcripts with multiple clusters in the 3’UTR, ΣB, ΔΣB and number of binding sites in a cluster represent values of the cluster closest to the PAS. Proximity to PAS in transcripts of multiple clusters represents the median pattern for the clusters (for example, in a UTR with 5 clusters, 4 of which distant to the PAS, the median was considered distant to the PAS). The empirical mean for each column of the data matrix was calculated (sample mean of each column, shifted to zero to center data). Data were centered and scaled and a covariance matrix for the seven features was calculated (**Extended Fig. 8a**). This covariance matrix was used to calculate eigenvectors and eigenvalues, as described ^63^. Eigenvalues were sorted in descending order and *K* largest eigenvalues were selected. *K* is the desired number of dimensions (Principal Components) of a new feature subspace Y with K ≤ n (*K* = 2 for **Extended Fig.8c** and *K* = 3 for **Extended Fig.8e**). A projection matrix (W) was created from the selected (*K*) eigenvalues through orthogonal transformation of the original dataset (*X*) in order to obtain a *K*-dimensional feature subspace Y. Proportion of variance, cumulative variance, factor loadings and eigenvalues explained by each component were recorded. Functional mRNA classes (**Extended Fig.8c**) and Dazl code groups (1 – 21, **Extended Fig.8e**) were identified and mapped onto the feature space (*Y*) by k-means clustering ^64^. PCA was conducted in R using the *prcomp()* function. To visualize subgrouping within functional mRNA classes (**Extended Data Fig.8d**), the Barnes-Hut t-SNE implementation in R ^65^ was used with the recommended parameters (perplexity 5 – 30, iterations 5 – 3000) as described ^66^.

### Derivation of the Dazl Code

Seven features of Dazl clusters and of transcripts with Dazl binding sites in 3’UTR (number of clusters in 3’UTR, ΣB, ΔΣB, number of binding sites in a cluster, UTR length, proximity to PAS, transcript level) were utilized to further group transcripts in each functional mRNA class (**Fig.4d**). In transcripts with multiple clusters in the 3’UTR, ΣB, ΔΣB and number of binding sites in a cluster represent values of the cluster closest to the PAS. Proximity to PAS in transcripts of multiple clusters represents the median pattern for the clusters (for example, in a UTR with 5 clusters, 4 of which distant to the PAS, the median was considered distant to the PAS). PCA and t-SNE independently identified 21 groups (1-21) in the 6 functional mRNA classes (**Extended Data Figs.7**,**8**). To create the Dazl code from identified groups 1-21, we first defined terciles (High, Median, Low) for each of the 7 features of Dazl binding patterns (number of clusters in 3’UTR, ΣB, ΔΣB, number of binding sites in a cluster, UTR length, proximity to PAS, transcript level) on the basis of significance testing (p < 0.05) for the deviation from the mean. The number of clusters of each tercile type (H, M or L) for each of the 7 features was then counted in each group. This yielded a data matrix with count of feature tercile (example: [group 1; ΣB]; H = 2, M = 27, L = 8, Total = 37 Clusters, **Extended Data Fig.8f**). The tercile count per feature (per group) was then normalized to total number of clusters in the group to obtain fraction of each feature tercile in a group (example: [group 1; ΣB]; H = 0.05, M = 0.73, L = 0.22, Total = 37 Clusters). For every group, the tercile for a feature that encompassed >50% of the clusters was utilized as the code for that group **(Extended Data Fig.8f)**.

### Linear Regression Analysis

Multiple regression analysis was performed with “dummy coding”, e.g. transformation of categorical variables into dichotomous variables ^67^. Variables in the Dazl-code were (N = 7): number of clusters in 3’UTR, ΣB, ΔΣB, number of binding sites in a cluster, UTR length, proximity to PAS, transcript level, expressed as terciles of their respective distributions.

Multiple regression was performed according to the general equation:

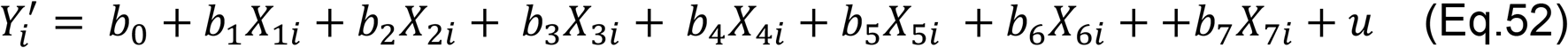

(Y’: predicted dependent, continuous variable (ΔRPF and ΔRNA), *b*_(i=0…7)_ : differential intercept linear coefficients, X_ni_: independent variables, u: error term). The differential intercept linear coefficients associated with each dummy variable are the expected difference in the mean of the outcome variable for that variable, compared to the reference group (MM class), with all other predictors constant ^67,68^. The final regression model was:

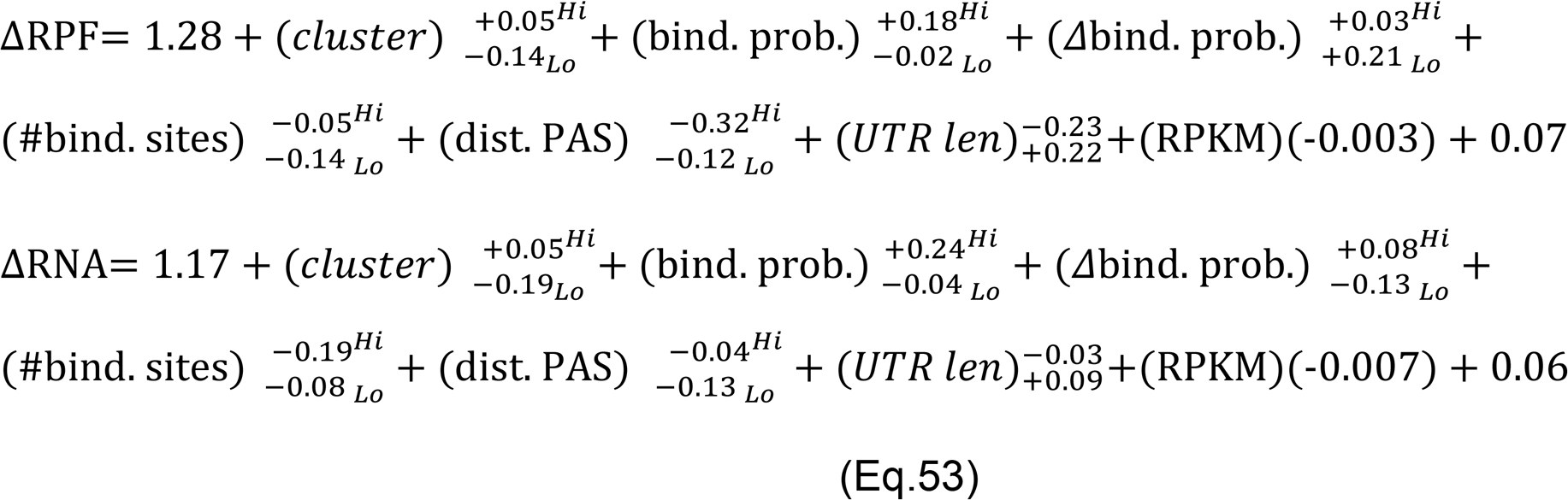

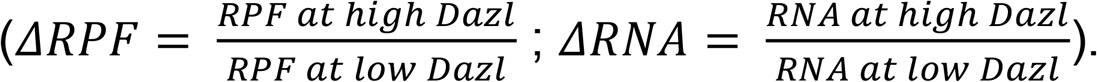

Regression analysis was performed using Scikit-learn ^69^ and Statsmodels ^70^ modules in Python 3.6.5.

### Decision Tree Classifier

We employed a Chi-squared Automatic Interaction Detection (CHAID) algorithm, which makes no assumption about underlying data ^71,72^, in order to determine how categorical independent variables (seven transcript and cluster features, above) best combine to predict the functional mRNA classes. A data matrix was formed using classes of Y (transcript and cluster features) as columns and categories of the predictor X (functional mRNA classes) as rows. The expected cell frequencies under the null hypothesis were estimated as described ^71^. The observed cell frequencies and the expected cell frequencies were then used to calculate Pearson chi-squared statistic, according to:

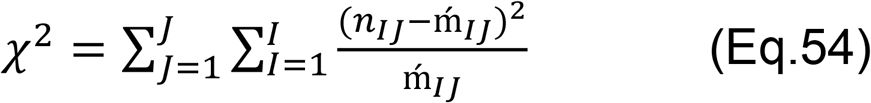

(*n_IJ_* is the observed cell frequency for cell *(x_n_* = *I* | *y_n_ = j)*. *m_IJ_* is the estimated expected cell frequency for cell *(x_n_* = *I* | *y_n_ = j)* from independence model ^71,72^.

The p value is:

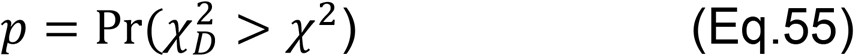

*X*_*D*_^*2*^ follows a Chi-squared distribution with degrees of freedom *d = (J − 1) (I − 1)*

Pr: probability. The adjusted p-value is calculated as Bonferroni multiplier ^73^.

CHAID analysis was performed using CHAID 5.3.0 (ref.^74^) in Python 3.6.5.

### Gene Ontology Analysis

GO term analyses for transcripts in groups 1-21 (**Fig.4d**) was performed with REACTOME (refs. ^75,76^) using a hypergeometric statistical test and Benjamini and Hochberg FDR correction (significance level of 0.05) to identify enriched terms after multiple testing correction ^77^. Redundant GO terms were merged to create a parent term. Transcripts for each Dazl group (1-21) were clustered using Ward’s minimum variance method in R ^78^ and plotted as a heatmap using ggplot2 ^48^ (**Fig.4d**).

### Pathway Analysis

Pathways (**Extended Data Fig.8h**) were obtained from REACTOME ^75,76^. mRNA classes were mapped on pathways with Cytoscape ^79^.

