## Supplementary Material for "The kinetic landscape of an RNA binding protein in cells"

**Supplementary Table S1 | Codon optimized *Mus musculus* Dazl (RRM) DNA construct** (amino acids 32 -117) **and primers for cloning.**

---

Dazl (RRM) DNA construct

SacI and XhoI restriction sites are underlined. Complete DNA construct was purchased from Genscript.

```
GGAAATATAGAGCTCTTGCCGGAAGGCAAGATCATGCCGAACACCGTATTCGTAGGAGGAATAG
ACGTACGCATGGACGAAACCGAAATCCGCTCTTTTTTCGCACGCTACGGCTCTGTAAAGGAGGT
TAAAATAATCACGGACAGAACGGGGGTTTCGAAAGGCTACGGATTCGTCTCTTTCTACAACGAT
GTTGACGTTCAGAAAATAGTAGAGTCTCAGATAAACTTTCATGGGAAGAAACTGAAGCTGGGCC
CGGCTATCCGCAAACAATAATGACCTCGAGGGCTGCAA
```

---

Primers for cloning

SacI and XhoI restriction sites are underlined.

Dazl Forward

5’-GGAAATATAGAGCTCTTGCCGGAAGGCAAGATCATGC

Dazl Reverse

5’-TTGCAGCCCTCGAGGTCATTATTGTTTGCGGATA

**Supplementary Table S2 | Sequencing adapters and primers.**

RNA linkers (Dharmacon)

RL5: 5’-OH AGG GAG GAC GAU GCG G 3’-OH

RL5D: 5’-OH AGG GAG GAC GAU GCG Gr(N)r(N) r(N)r(N)G 3’-OH

RL3: 5’-P GUG UCA GUC ACU UCC AGC GG 3’-puromycin

DNA primers (Operon)

DP5: 5’-AGG GAG GAC GAT GCG G-3’

DP3: 5’-CCG CTG GAA GTG ACT GAC AC-3’

Solexa Fusion Primers (Operon)

SSP1: 5'-CTA TGG ATA CTT AGT CAG GGA GGA CGA TGC GG-3'

Circularization RT primer (Dharmacon)

5’Phos/(GGTTA)(CCGCATCGTCCTCCCT)(CCCTATAGTGAGTCGTATTA)/iSp18/CACTCA/iSp18/(CCGCTGGAA GTGACTGACAC)3’

Antisense DP5 Antisense T7 Promoter DP3

1) 5’Phos-GNNNN CGTGAT CCGCATCGTCCTCCCTC CCTATAGTGAGTCGTATTA - iSp18 - CACTCA -iSp18 – CCGCTGGAAGTGACTGACAC

2) 5’Phos-GNNNN ACATCG CCGCATCGTCCTCCCTC CCTATAGTGAGTCGTATTA - iSp18 - CACTCA -iSp18 – CCGCTGGAAGTGACTGACAC

3) 5’Phos-GNNNN GCCCTA CCGCATCGTCCTCCCTC CCTATAGTGAGTCGTATTA - iSp18 - CACTCA -iSp18 – CCGCTGGAAGTGACTGACAC

4) 5’Phos-GNNNN TGGTCA CCGCATCGTCCTCCCTC CCTATAGTGAGTCGTATTA - iSp18 - CACTCA -iSp18 – CCGCTGGAAGTGACTGACAC

5) 5’Phos-GNNNN CACAGT CCGCATCGTCCTCCCTC CCTATAGTGAGTCGTATTA - iSp18 - CACTCA -iSp18 – CCGCTGGAAGTGACTGACAC

6) 5’Phos-GNNNN ATTGGC CCGCATCGTCCTCCCTC CCTATAGTGAGTCGTATTA - iSp18 - CACTCA -iSp18 – CCGCTGGAAGTGACTGACAC

Complementary barcode sequence

1) ATCACGNNNNG……………

2) CGATGTNNNNG……………

3) TAGGGCNNNNG…………..

4) TGACCANNNNG…………..

5) ACTGTGNNNNG…………..

6) GCCAATNNNNG…………..

| Time (s) | Dazl: 4.2x L: 2.6 mW | Dazl: 4.2x L: 1 mW | Dazl: 1x L: 2.6 mW | Dazl: 1x L: 1 mW | Stratalinker |
|---|---|---|---|---|---|
| 0 | $5 \cdot 10^6$ | $6 \cdot 10^6$ | $4 \cdot 10^6$ | $3 \cdot 10^6$ | $5 \cdot 10^6$ |
| 30 | $3 \cdot 10^6$ | $3.6 \cdot 10^6$ | $4 \cdot 10^6$ | $8 \cdot 10^6$ | $5 \cdot 10^6$ |
| 180 | $1.9 \cdot 10^6$ | $2.4 \cdot 10^6$ | $4 \cdot 10^6$ | $5 \cdot 10^6$ | $5 \cdot 10^6$ |
| 680 | $0.6 \cdot 10^6$ | $1.2 \cdot 10^6$ | $2 \cdot 10^6$ | $3 \cdot 10^6$ | $5 \cdot 10^6$ |

**Supplementary Table S3 | Number of cells used in each crosslinking experiment**
(L: laser power)

| Time (s) | Dazl: 4.2x<br>L: 2.6 mW | Dazl: 4.2x<br>L: 1 mW | Dazl: 1x<br>L: 2.6 mW | Dazl: 1x<br>L: 1 mW | Stratalinker |
|---|---|---|---|---|---|
| 30 | 88% | 98% | 80% | 91% | 91% |
| 180 | 79% | 92% | 82% | 87% | 84% |
| 680 | 87% | 81% | 93% | 91% | 83% |

**Supplementary Table S4 | Cell Viability after each crosslinking experiment**
(L: laser power). Cell viability was measured by Trypan-blue staining and cell counting in a hemocytometer (Materials and Methods).

| Conditions | 680 s | 180 s | 30 s | 0 | 680 s | 180 s | 30 s | 0 | 680 s | 180 s | 30 s | 0 | 680 s | 180 s | 30 s | 0 |
|---|---|---|---|---|---|---|---|---|---|---|---|---|---|---|---|---|
| | Dazl: 4.2x | | | | Dazl: 1x | | | | Dazl: 4.2x | | | | Dazl: 1x | | | |
| | Laser: 2.6 mW | | | | Laser: 2.6 mW | | | | Laser: 1 mW | | | | Laser: 1 mW | | | |
| Post processed reads [(a)] | 3,372,238 | 466,053 | 357,206 | 13,800 | 545,542 | 283,506 | 150,313 | 12,720 | 249,005 | 364,176 | 141,804 | 15,650 | 394,016 | 227,026 | 175,420 | 8,730 |
| Mapped Reads [(b)] | 1,140,415 | 341,785 | 214,324 | 828 | 256,405 | 172,939 | 111,232 | 865 | 186,754 | 185,730 | 90,755 | 1,001 | 165,487 | 154,378 | 112,269 | 567 |
| % Reads Mapped | 33.81 | 73.33 | 60.00 | 6.0 | 47.00 | 61.00 | 74.00 | 6.8 | 75.00 | 51.00 | 64.00 | 6.4 | 42.00 | 68.00 | 64.00 | 6.5 |
| Correction factor [(c)] | 0.89 | 2.28 | 2.56 | 1 | 2.88 | 2.22 | 3.11 | 1 | 1.84 | 2.17 | 2.2 | 1 | 1.88 | 2.54 | 3 | 1 |
| Reads - Peak Intersection [(d)] | 252,932 | 185,659 | 173,943 | 0 | 204,474 | 86,071 | 92,228 | 0 | 153,860 | 48,334 | 74,552 | 0 | 79,527 | 11,271 | 14,910 | 0 |

**Supplementary Table S5 | Sequencing and read processing statistics.**

[(a)] Post processed reads: Reads remaining after de-multiplexing, adapter removal and PCR duplicate collapsing.

[(b)] Mapped reads: Reads mapped to mouse genome (mm10).

[(c)] Correction factor: Intensity per read obtained by normalizing number of reads per condition with total crosslinked RNA.

[(d)] Reads-Peak intersection: Number of reads corresponding to Dazl binding site peaks common to all KIN-CLIP conditions.

| Conditions | 680 s | 180 s | 30 s | 0 | 680 s | 180 s | 30 s | 0 | 680 s | 180 s | 30 s | 0 | 680 s | 180 s | 30 s | 0 |
|---|---|---|---|---|---|---|---|---|---|---|---|---|---|---|---|---|
| | Dazl: 4.2x | | | | Dazl: 1x | | | | Dazl: 4.2x | | | | Dazl: 1x | | | |
| | Laser: 2.6 mW | | | | Laser: 2.6 mW | | | | Laser: 1 mW | | | | Laser: 1 mW | | | |
| Bulk Crosslinking Intensity ($10^6$) | 1.012 | 0.775 | 0.537 | $10^{-5}$ | 0.722 | 0.384 | 0.346 | $10^{-5}$ | 0.343 | 0.403 | 0.199 | $10^{-5}$ | 0.311 | 0.392 | 0.336 | $10^{-5}$ |

**Supplementary Table S6 | Bulk crosslinking intensity for each crosslinking condition.**

Bulk crosslinking (AU; pixel density as described in Image J) was measured as described in Materials and Methods.

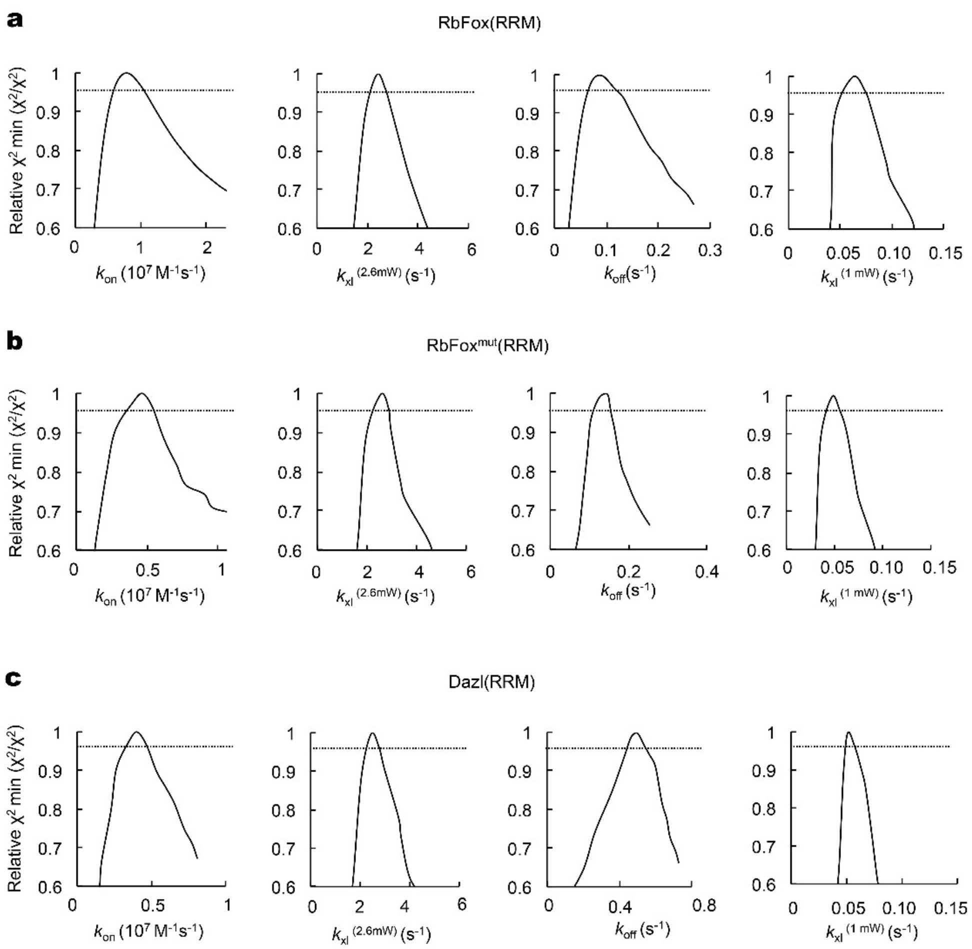


**Supplementary Figure S1 | fs laser crosslinking fit space parameters.**

Fit space analysis (KINTEK) for obtained kinetic parameters ($k_{on}$, $k_{xl}^{2.6mW}$, $k_{off}$ and $k_{xl}^{1mW}$) for **(a).** RbFox(RRM), **(b)** RbFox$^{Mut}$(RRM) and **(c)** Dazl(RRM). (**Fig.1e**). The relative $X^2$ represents the smallest (optimal) $X^2$ divided by the $X^2$ obtained for the entire thermodynamic model. For the optimal parameter value, the relative $X^2$ = 1. Horizontal lines mark the 95% confidence interval.

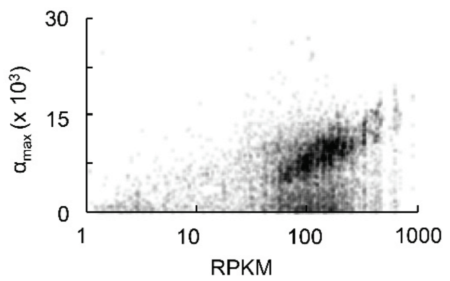


**Supplementary Figure S2 | Determination of fractional occupancy ($\Phi^{max}$)**

Maximal amplitude ($\alpha^{max}$: probability of Dazl bound to the fraction of a given binding site that is accessible during the course of the experiment, extrapolated to saturating concentrations of Dazl) plotted vs. level of the corresponding transcript (RPKM). Eq.44 (Materials and Methods) is used to calculate the maximal fractional occupancy ($\Phi^{max}$).

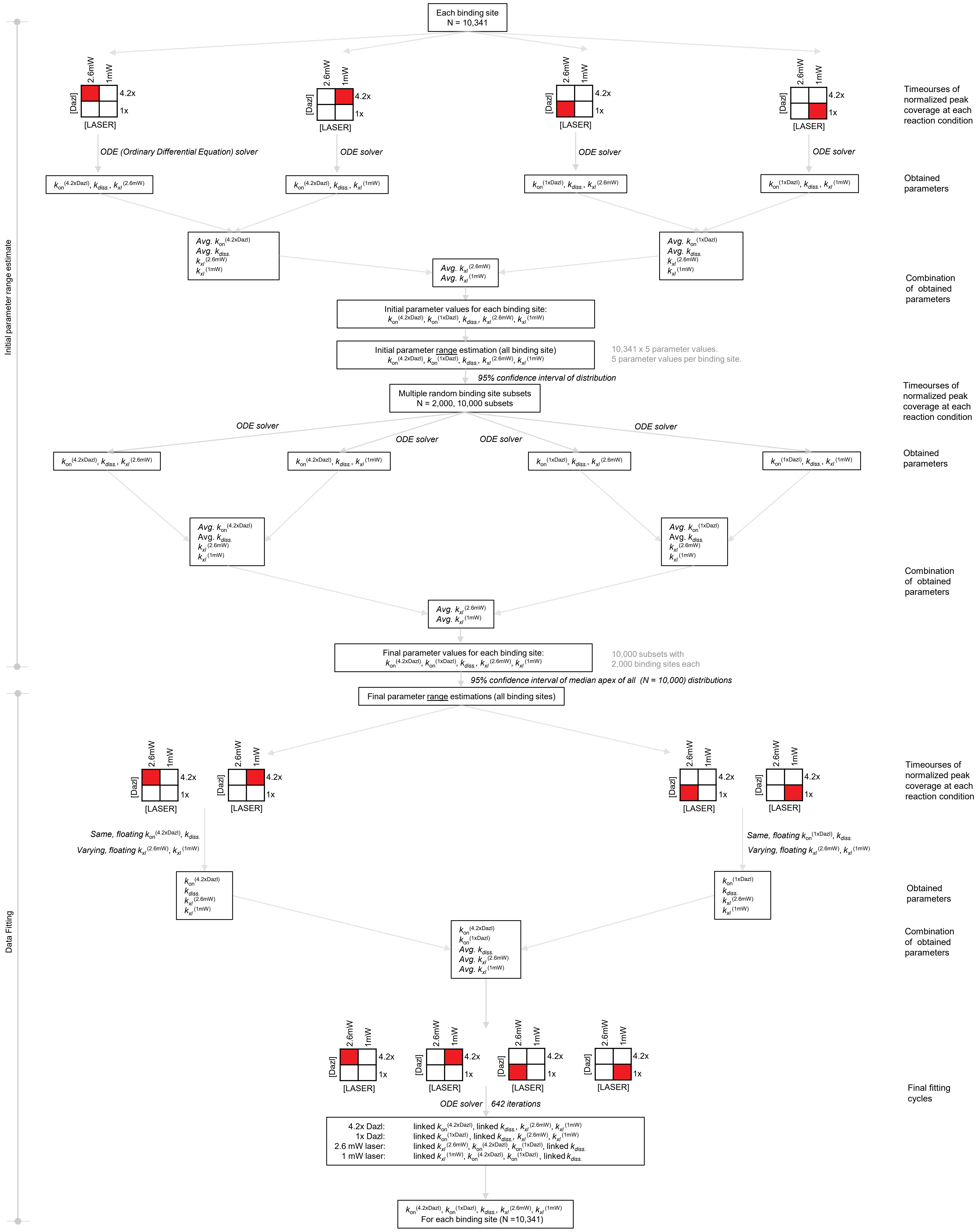


**Supplementary Scheme S1 | Numerical data fitting process**

Steps for the numerical fitting of crosslinking timecourses to calculate kinetic parameters. Square boxes represent KIN-CLIP conditions (red).

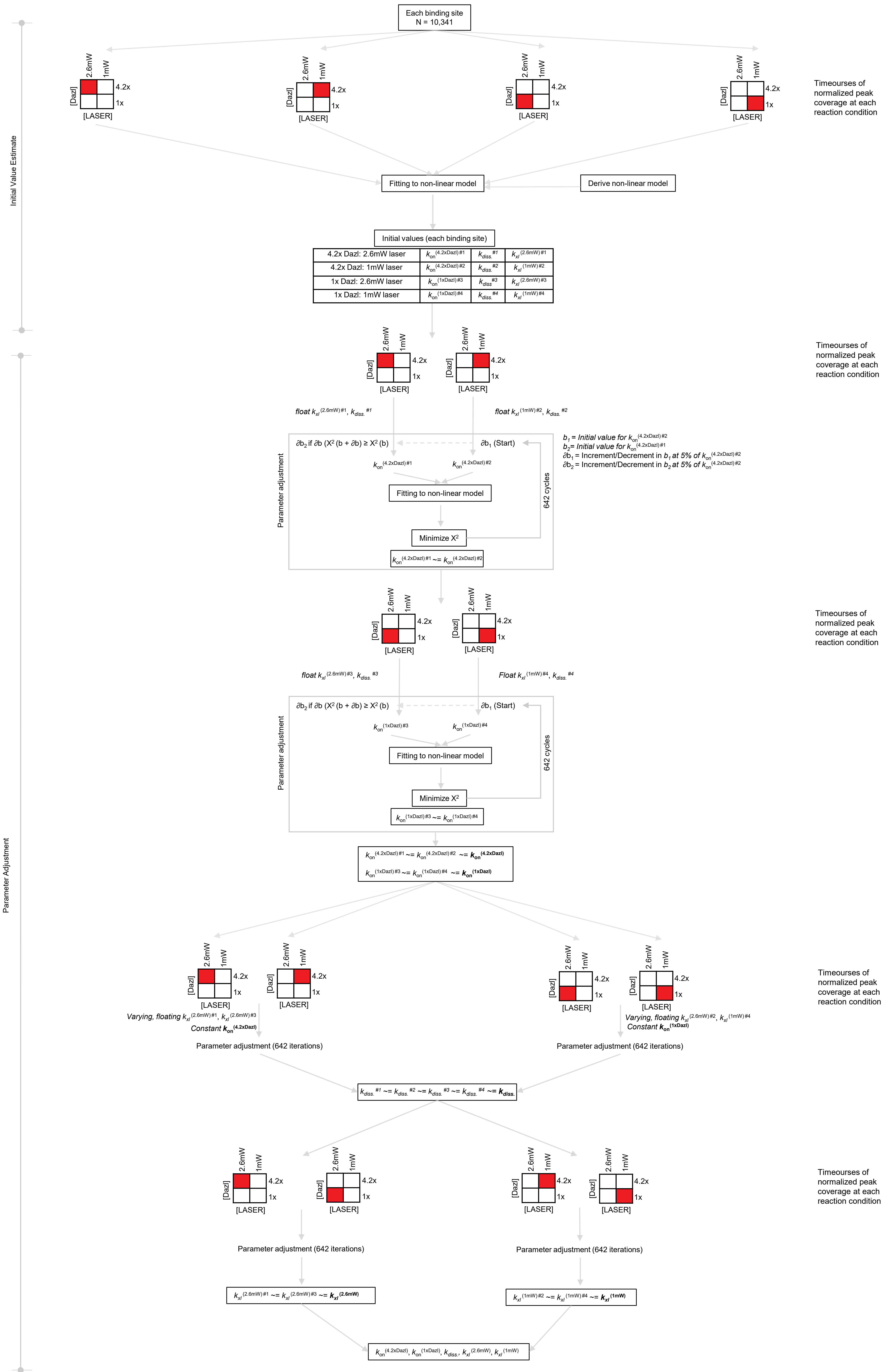


**Supplementary Scheme S2 | Analytical data fitting process**

Steps for the numerical fitting of crosslinking timecourses to calculate kinetic parameters. Square boxes represent KIN-CLIP conditions (red).
